# RAI1 Regulates Activity-Dependent Nascent Transcription and Synaptic Scaling

**DOI:** 10.1101/523456

**Authors:** Patricia M. Garay, Alex Chen, Takao Tsukahara, Rafi Kohen, J. Christian Althaus, Margarete A. Wallner, Roman J. Giger, Michael A. Sutton, Shigeki Iwase

**Affiliations:** Neuroscience Graduate Program, The University of Michigan Medical School, Ann Arbor, MI 48109, USA; Department of Molecular & Integrative Physiology, University of Michigan, Ann Arbor, Michigan 48109, USA; Molecular and Behavioral Neuroscience Institute, University of Michigan, Ann Arbor, Michigan 48109, USA; Department of Human Genetics, University of Michigan, Ann Arbor, Michigan 48109, USA; College of Literature, Science, & the Arts, University of Michigan, Ann Arbor, MI 48109, USA; Department of Cell and Developmental Biology, University of Michigan School of Medicine, Ann Arbor, MI, United States; Cellular and Molecular Biology Graduate Program, University of Michigan Medical School, Ann Arbor, MI, United States; Department of Neurology, University of Michigan Medical School, Ann Arbor, MI, United States

## Abstract

Long-lasting forms of synaptic plasticity such as synaptic scaling are critically dependent on transcription. Activity-dependent transcriptional dynamics in neurons, however, have not been fully characterized, because most previous efforts relied on measurement of steady-state mRNAs. Here, we profiled transcriptional dynamics of primary neuronal cultures undergoing network activity shifts using nascent RNA sequencing. We found pervasive transcriptional changes, in which ~45% of expressed genes respond to network activity shifts. Notably, the majority of these genes respond to increases or decreases of network activity uniquely, rather than reciprocally. We further linked the chromatin regulator Retinoic acid induced 1 (RAI1), the Smith-Magenis Syndrome gene, to the specific transcriptional program driven by reduced network activity. Finally, we show that RAI1 is essential for homeostatic synaptic upscaling but not downscaling. These results demonstrate the utility of *bona fide* transcription profiling to discover mechanisms of activity-dependent chromatin remodeling that underlie normal and pathological synaptic plasticity.

## Introduction

Proper cognitive development and brain function relies on synaptic plasticity – the ability of synapses to strengthen or weaken in response to sensory or neuromodulatory inputs. Synaptic scaling is one mechanism of plasticity, which buffers potentially destabilizing patterns of network activity (Abbott and Nelson, 2000; Miller and MacKay, 1994; Turrigiano, 2008). In response to a sustained increase in neuronal firing rate, neurons decrease, or “scale-down”, the receptivity of the neuron to excitatory neurotransmitters. Conversely, global decreases in firing rate causes neurons to “scale-up” and increase synaptic efficacy. Synaptic scaling is thought to accommodate other forms of plasticity, such as long-term potentiation (LTP), that impose long-lasting increase of individual synaptic efficacy, which if left uncompensated, would result in circuits that are overly active (Turrigiano, 2017). Synaptic scaling appears to play important roles in neurodevelopment, learning, and memory (Fernandes and Carvalho, 2016; Yee et al., 2017). Dysregulated homeostatic plasticity is a common pathological hallmark in several neurodevelopmental disorders, including Fragile X (Soden and Chen, 2010), Rett syndrome (Zhong et al., 2012), tuberous sclerosis (Bateup et al., 2013), Kleefstra syndrome (Benevento et al., 2016), and has been proposed to underlie autism spectrum conditions (Bourgeron, 2015). To understand how homeostatic plasticity contributes to normal and pathological brain development, identifying the molecular mechanisms underlying synaptic scaling is an important first step.

Long-lasting forms of synaptic plasticity, including synaptic scaling, require *de novo* synthesis of RNAs, which in turn produce the proteins that directly modulate synaptic efficacy (Benito and Barco, 2015; Ibata et al., 2008; Igaz et al., 2002). Increased neuronal firing leads to classic signal transduction cascades that eventually phosphorylate key transcription factors (TFs) such as cyclic AMP-response binding protein (West et al., 2002). TFs bind to specific DNA sequences, e.g. cyclic AMP-response element, and play essential roles in expression of immediate early genes (IEG), which encode key players in synaptic scaling, such as ARC (Bramham et al., 2008) and HOMER1 (Brakeman et al., 1997). Reduction of network activity leads to downregulation of these IEGs, while inducing the expression of genes that can scale up net synaptic efficacy (Schaukowitch et al., 2017). The gene expression programs triggered by reductions in network activity involves SRF, another key TF, which cooperates with the transcriptional coactivator ELK1 (Schaukowitch et al., 2017). However, molecular mechanisms underlying the transcriptional response to activity shifts remain incompletely understood.

In order to control DNA accessibility of the transcription machinery, i.e. RNA polymerase II and associated factors, TFs need to collaborate with chromatin regulators. Chromatin regulators can deposit or remove a variety of modifications on DNA and histones, thereby influencing higher order chromatin structure. The strong linkage between cognitive disorders and chromatin-regulatory genes suggests that activity-dependent chromatin reorganization is essential for proper brain development and mental health (Ebert and Greenberg, 2013; Guzman-Karlsson et al., 2014; Mullins et al., 2016). The roles of many chromatin regulators have been described in activity-dependent gene expression that are linked to LTP and memory (Alarcón et al., 2004; Bourtchouladze et al., 2003; Guan et al., 2009; Gupta-Agarwal et al., 2012; Gupta-Agarwal et al., 2014; Iwase et al., 2016; Kerimoglu et al., 2013; Lim et al., 2017; Neelamegam et al., 2012; Oike et al., 1999; Oliveira, 2016; Rudenko et al., 2013; Vogel-Ciernia et al., 2013; Wang et al., 2015). However, only a handful of chromatin regulators, i.e., TET3 DNA demethylase (Yu et al., 2015), EHMT1/2 histone H3K9 methyltransferases (Benevento et al., 2016), and L3MBTL1 methyl-histone binding factor (Mao et al., 2018) have been well-characterized for activity-dependent chromatin remodeling that underlies synaptic scaling. These three chromatin regulators participate distinctly in synaptic scaling. TET3 is essential for both up-and downscaling, whereas L3MBTLl1 is essential only for synaptic downscaling.

In order to dissect the mechanism by which chromatin regulators impart distinct effects on synaptic scaling, it is crucial to carefully monitor bidirectional transcriptional responses to increases or decreases in network activity. Most studies have addressed this issue using quantification of steady-state mRNA levels, using RT-qPCR, cDNA microarray, and mRNA-seq. The brain is characterized by its notorious complexity of post-transcriptional regulation, including activity-dependent mRNA splicing (Hermey et al., 2017), mRNAs stability (Widagdo and Anggono, 2018), mRNA transport and local translation (Glock et al., 2017). Therefore, changes in steady-state mRNA levels do not necessarily indicate a direct transcriptional impact of a given chromatin regulator. Thus, reliance on steady-state mRNA measurements may obscure the roles of chromatin regulators in transcription.

In the present work, we developed genome-wide measurement of *bona fide* transcriptional dynamics in response to bidirectional network activity alterations. We then used this approach to uncover a novel role for the chromatin regulator Retinoic acid induced 1 (RAI1) in the transcriptional program specifically elicited by reduced neural activity. RAI1 is a nucleosome binding protein (Darvekar et al., 2012; Darvekar et al., 2013) that is consistently expressed in the brain during neurogenesis and throughout adulthood, in both mice and humans (Huang et al., 2016). *RAI1* is associated with two human intellectual disability syndromes. *RAI1* haploinsufficiency leads to Smith-Magenis Syndrome (SMS, MIM: 182290), while duplication of the genomic region containing *RAI1* results in Potocki-Lupski syndrome (PTLS, MIM: 610883) (Bi et al., 2004; Girirajan et al., 2005; Potocki et al., 2007; Slager et al., 2003), Heterozygous and homozygous Rai1-knockout (KO) mice exhibit many of the symptoms of SMS patients including learning deficits, abnormal circadian and social behavior, as well as obesity (Bi et al., 2005; Bi et al., 2007; Lacaria et al., 2013) (Huang et al., 2018; Huang et al., 2016). Furthermore, heterozygous *Rai1*-KO mice display altered gene expression profiles and reduced dendritic spine density in the prefrontal cortex. These studies implicate a role for RAI1 in gene expression, neuronal structure, and behavior, but the precise role for RAI1 in activity-dependent transcription and synaptic plasticity remains unclear. Here, using nascent RNA sequencing to monitor *bona fide* transcriptional events during network activity shifts, we define a specific role for RAI1 in the transcription program elicited by reduced network activity and show further that RAI1 is essential for homeostatic upscaling during chronic activity suppression.

## Results

### Altered neuronal network-activity triggers genome-wide transcriptional changes

We first sought to develop an experimental paradigm that allows us to monitor *bona fide* transcriptional dynamics in neuronal networks. To overcome the major limitation of conventional RNA-seq. i.e. profiling only steady-state RNA quantities, we adopted Bromouridine-sequencing (Bru-seq), a genome-wide profiling technique of nascent transcripts (Paulsen et al., 2014; Paulsen et al., 2013). We prepared primary forebrain neuronal cultures from E18 mouse embryos, and allowed them to mature for 17 days *in vitro* (DIV). To monitor bidirectional transcriptional responses to activity shifts, network activity was either elevated by 20 μm bicuculline (BIC, a GABA_A_-receptor antagonist) or suppressed by 1 μm tetrodotoxin (TTX, a sodium channel blocker) for 4 hours. During the last 20 minutes of BIC or TTX treatment, bromouridine (BrU) was added to the culture medium to label newly-synthesized transcripts. The labeled RNAs were isolated by immunoprecipitation using an anti-BrU antibody and subjected to next-generation sequencing (Fig. 1A).

**Figure 1.**
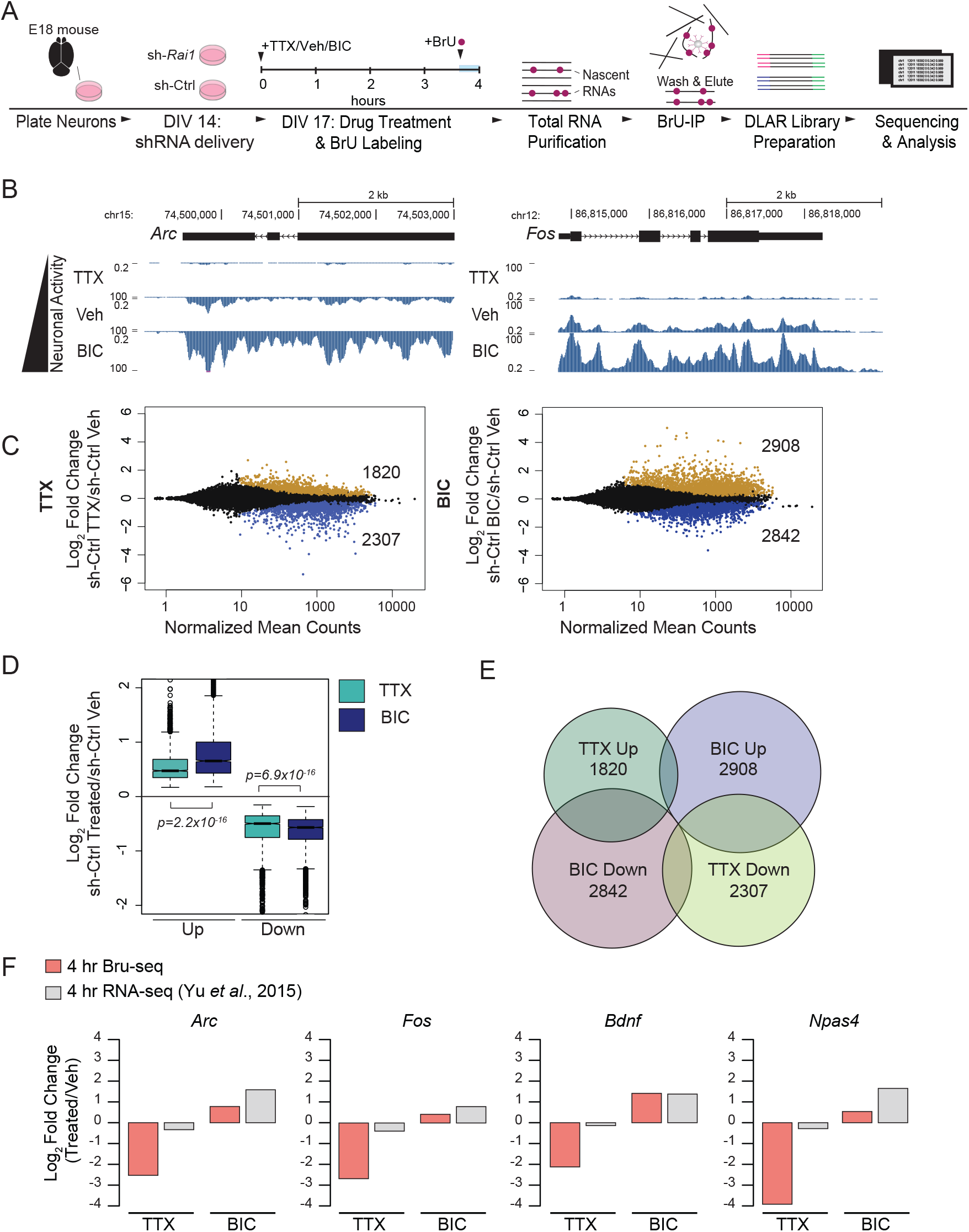
Genome-wide transcriptional response to bi-directional activity alterations. **(A)** Experimental procedure. **(B)** UCSC Browser views of Bru-seq signals at *Arc* and *Fos.* Intronic reads are characteristic of nascent RNA. **(C)** Differential gene expression analysis (DESeq2) reveals widespread transcriptional changes in response to TTX and BIC (p_adj_<0.05). **(D)** BIC-response genes show a greater median fold change (Wilcoxon rank-sum test, upregulated genes: p=2.2 x 10^-16^, downregulated genes: p=6.9 x 10^-16^). Whiskers represent 1.5 times the inter-quartile range (IQR) and the notch represents the 95% confidence interval of the median. **(E)** The majority of TTX and BIC response genes are uniquely regulated (70%). 24% of genes are reciprocally regulated and 6% are commonly regulated. **(F)** Downregulation of immediate early gene in the TTX condition is captured more sensitively in Bru-seq data compared to mRNA-seq data (Yu et al., 2015).

We first validated the results of Bru-seq by examining transcription of known activity-dependent genes individually. As shown in Figure 1B, we found the expected changes of *Arc* and *Fos*, which were downregulated by TTX and upregulated by BIC. Abundant intronic reads indicate that detected transcripts were recently generated and yet to be spliced. Other well-characterized activity-dependent genes such as *Npas4, Egr1, Homer1, Tet3*, and *Txnip* showed expected transcriptional dynamics (Table S1). Thus, Bru-seq reliably captures known transcriptional responses to bidirectional shifts in network activity.

Following individual gene validation, we characterized genome-wide transcriptional responses to BIC and TTX. Differential gene expression analysis of the Bru-seq data using DESeq2 (Love et al., 2014) revealed widespread transcriptional changes, in which 45% of expressed genes (7,592/16,682) were significantly up- or down-regulated by network activity shifts (p_adj_ < 0.05, Fig. 1C). BIC increased transcription of 2,908 genes, whereas TTX did so for 1,820 genes. The magnitude of transcriptional induction is higher in BIC treatment compared to TTX (Fig. 1D). Meanwhile, a similar number of genes were transcriptionally suppressed upon BIC (2,842) and TTX (2,307) treatments. A relatively small fraction of genes, 24% (1,798 of 7,592 activity-response genes), displayed reciprocal changes between BIC and TTX treatments, e.g. upregulation by BIC and downregulation by TTX (Fig. 1E). An even smaller fraction, 6% (487/7,592) of activity-response genes altered their transcription levels in the same direction after BIC and TTX treatments. The remaining 70% of genes (5,307/7,592) responded to BIC or TTX uniquely. Supplemental Table 1 lists genes that displayed greater than 2-fold changes in transcription upon network-activity shifts.

We next analyzed two published mRNA-seq datasets, which profiled the steady-state transcriptome of mouse cortical neurons treated with TTX or BIC for 4 or 6 hours (Schaukowitch et al., 2017; Yu et al., 2015) with identical processing and analyses (see methods). We found that Bru-seq identified a greater number of differentially-expressed (DE) genes compared to conventional mRNA-seq. The DE genes found in each dataset only partially overlapped (Fig. S1A). However, the groups of DE genes identified in the mRNA-seq dataset of 4-hour treatments were shifted in the same direction in Bru-seq (Fig. S1B). In contrast, the group of DE genes of the 6-hour RNA-seq dataset did not show any noticeable transcriptional changes in the Bru-seq data (Fig. S1C). These results suggest that transcriptional responses are highly dynamic and correlate with steady-state mRNA levels only in a narrow window of time (< 2 hr).

Conventional mRNA-seq may not be the ideal approach to detect downregulation of transcription, because after transcription ceases, synthesized RNAs persist for certain periods of time. To test whether Bru-seq can detect transcriptional downregulation sensitively, we compared the induction and suppression of known immediate-early genes in our Bru-seq and published mRNA-seq datasets. We found significantly larger suppression of *Fos, Arc, Bdnf*, and *Npas4* (4-to 16-fold) by TTX treatment in Bru-seq compared to conventional mRNA-seq, in which downregulation was less than 2-fold (Fig. 1F). In the Bru-seq data, the four genes showed smaller magnitudes of upregulation in response to BIC, likely because the early transcriptional induction is largely complete 4 hours after BIC treatment (Fig. S1C). These data highlight an advantage of the Bru-seq approach to probe mechanisms underlying highly-dynamic activity-dependent transcription.

We next examined the cell-type specificity of activity-dependent genes in our datasets. Recent studies have reported that different cell types such as astrocytes and neuronal subtypes induce distinct sets of genes in an activity-dependent manner (Hasel et al., 2017; Hrvatin et al., 2018). Using immunocytochemistry of a set of well-established markers, NeuN, GAD67, GFAP, CD11b, and Olig2, we estimated that our cultures consist of 41% excitatory neurons, 11% inhibitory neurons, 33% astrocytes, 15% of cells within the oligodendrocyte lineage, and no microglia (Fig. S2A-B). Indeed, several non-neuronal genes are represented in our dataset, including *Thbs1*, a known synaptic regulator specifically expressed in astrocytes (Risher and Eroglu, 2012). Gene ontology analysis of the Bru-seq data detected enrichment of biological processes specific for both neurons and non-neuronal cell types (Table S2). For example, regulation of axon diameter (p_adj_ = 2.6 x 10^-5^), axonal transport of mitochondrion (p_adj_ =3.5 x 10^-4^), and glial cell proliferation (p_adj_: 7.8 x 10^-3^) represent genes that are transcriptionally induced upon BIC treatment, while transcription of genes involved in astrocyte activation are down-regulated by BIC (p_adj_: 9.8×10^-5^). TTX treatment leads to increased transcription of myelin maintenance genes (p_adj_: 7.4×10^-3^). Reciprocally enriched biological processes include interneuron migration (BIC Down, p_adj_: 1.7×10^-3^, TTX Up, p_adj_: 7.0×10^-2^) and neuropeptide signaling pathway (BIC Up, p_adj_: 8.5×10^-7^, TTX down, p_adj_: 4.7×10^-5^). When we intersected published cell type-enriched genes (Zhang et al., 2014) (see Methods) with the Bru-seq data, indeed, some neuronal- and non-neuronal genes transcriptionally respond to activity shifts (Fig. S2C).

### RAI1 suppresses the TTX-induced transcriptional program in resting network

The superior sensitivity of Bru-seq over conventional RNA-seq prompted us to assess the role of RAI1 in activity-dependent gene expression. We first determined RAI1 expression in our culture systems. Publically-available databases indicate ubiquitous *Rai1* expression in a broad array of cell types in the brain (Fig. S3). Previous studies have demonstrated that *Rai1* mRNA expression rises from E13.5 to peak at P7, and its expression continues throughout adulthood across brain regions (Fragoso et al., 2014; Huang et al., 2016). We developed an anti-RAI1 antibody and confirmed that RAI1 protein was expressed in virtually all MAP2-positive neurons and primarily localized to the nucleus with subtle but detectable extra-nuclear signals in the soma (Fig. S4A). We found relatively low RAI1 levels in non-neuronal (MAP2-negative) cell nuclei (Fig. S4A). To examine if RAI1 protein levels or nuclear localization is altered by activity, we probed for RAI1 in cortical neurons treated with BIC, TTX, or Vehicle for 15 min, 1 hr, 2 hr, 4 hr, 8 hr, or 24 hr, using Western blots and fluorescent microscopy. Neither RAI1 protein levels nor localization were visibly altered in response to drug treatment at any time-point (Fig. S4 and S5). RAI1 has been shown to occupy a large fraction of active promoters in the mouse adult cortex (Huang et al., 2016). To examine whether RAI1 preferentially occupies activity-response genes, we utilized the published RAI1 chromatin immunoprecipitation sequencing (ChIP-seq) data obtained from the cortices of 8-week old mice (Huang et al., 2016). RAI1 ChIP-seq peaks were found at promoters of ~80% genes that are expressed at detectable levels, regardless of their activity-dependent transcriptional changes (Fig. S6A). We did not find any statistically-significant enrichment or depletion of RAI1 occupancy of TTX-or BIC-response genes (Fig. S6A). Thus, these data indicate that neuronal activity does not influence RAI1’s expression level or subcellular localization and that steady-state chromatin occupancy by RAI1 is not selective between BIC or TTX-response genes.

To directly test RAI1’s role in activity-dependent transcription, we went on to perform *Rai1* knockdown (KD) in the primary cultures using lentiviral vectors (LV) carrying *Rai1-* or scramble shRNAs (sh-Ctrl). To minimize impact of RAI1 loss on network connectivity, we delivered LV shRNA at DIV14, a time by which functional synapses have formed. Near complete loss of RAI1 protein was achieved by 3 days post-LV infection (Fig. 2A). Next, we modulated network activity of LV-treated cultures by applying TTX or BIC for four hours. Genome-wide transcription events were assessed by Bru-seq as described above.

**Figure 2.**
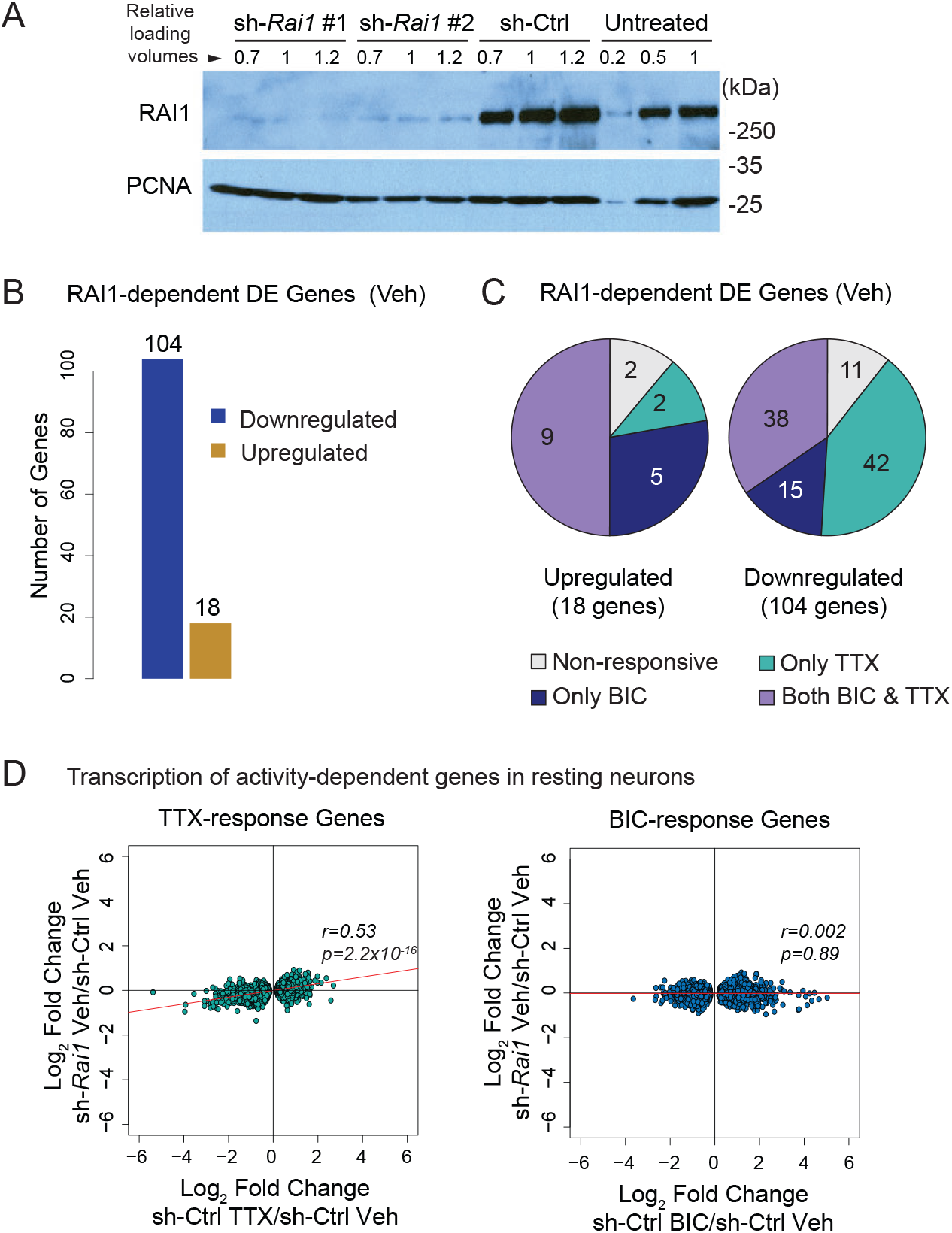
*Rai1-KD* alters transcription of TTX-response genes at the baseline. **(A)** Validation of *Rai1*-KD with Western blot. Mouse forebrain neuron cultures were transduced with lentivirus expressing sh-Rai1 or sh-Ctrl for three days. **(B)** Number of DESeq2-called differentially expressed genes (sh-Ctrl v sh-Rai1, p_adj_<0.05) after Vehicle treatment. **(C)** Many *Rai1*-KD DE-genes are TTX and BIC-response genes. **(D)** The fold changes of TTX- and BIC-response genes by *Rai1*-KD at baseline. Note that *Rai1*-KD cultures displays transcriptional profile similar to TTX-treated normal cultures. r= Spearman’s rank correlation coefficient.

We initially sought to establish if *Rai1*-KD alone was sufficient to alter nascent transcription in resting neuronal cultures. DESeq2 analysis revealed that 104 genes were downregulated and 18 genes were upregulated by *Rai1*-KD in the Vehicle-treated condition (p_adj_ <0.05, Fig. 2B, Table S3). The greater number of downregulated genes is consistent with previous studies demonstrating that RAI1 functions predominately as a transcriptional activator (Burns et al., 2010; Carmona-Mora et al., 2012; Carmona-Mora et al., 2010; Elsea and Williams, 2011; Girirajan et al., 2009; Huang et al., 2016). A majority of genes altered by *Rai1*-KD at baseline were either BIC- or TTX-response genes (Fig. 2C). Of note, however, BIC- or TTX-response genes were not significantly enriched in RAI1-dependent genes when gene groups were corrected for their expression levels (data not shown).

To further characterize the relationship between RAI1 deficiency and BIC- or TTX-response genes, we examined how individual genes behave upon *Rai1*-KD. We found a clear positive correlation between the normal transcriptional response to TTX and the transcriptional impairment by *Rai1*-KD at baseline (r=0.53, p=2.2×10^-16^, Spearman rank correlation coefficient, t-test, Fig. 2D, left panel). No correlation was found in BIC-response genes and *Rai1*-KD (Fig. 2D, right panel). The group of genes that respond reciprocally to TTX and BIC (Fig. 1E) showed similar correlation with all TTX-responsive genes (Fig. S6B). When we removed all DE genes upon *Rai1*-KD from the plot, the correlation remained significant (r=0.52, p=2.2×10^-16^, Fig. S6C), suggesting that the correlation was not solely driven by the DE genes. We also analyzed the published mRNA-seq of the *Rai1*-KO cortices (Huang et al., 2016) and found a similar trend in expression pattern of the TTX- and BIC-response genes (Fig. S6D). These data indicate that RAI1 deficiency shifts the transcriptional profile towards the TTX-treated state without drug application and that *Rai1*-KD does not impact transcription of non-reciprocal BIC-responsive genes.

### RAI1 deficiency promotes synaptic upscaling

Chronic perturbation of neuronal activity by BIC or TTX is known to induce decreases and increases in synaptic strength, which respectively, underlie homeostatic synaptic downscaling and upscaling (Abbott and Nelson, 2000; Miller and MacKay, 1994; Turrigiano, 2008). Given that *Rai1*-KD shifted the nascent transcriptome towards the TTX-like state, we next asked whether *Rai1*-KD would similarly shift excitatory synapse function towards a state similar to synaptic upscaling. We used sparse transfection of DIV12-14 hippocampal cultures with either *Rai1-* or scrambled shRNA, and recorded miniature excitatory postsynaptic currents (mEPSCs) from transfected pyramidal-like neurons 48 hours later. If *Rai1*-KD induces synaptic strengthening in a cell-autonomous manner, we would expect to see a rightward shift in the distribution of mEPSC amplitudes as is observed during synaptic upscaling following chronic activity suppression with TTX. Consistent with this idea, we found that expression of two distinct shRNAs targeting *Rai1* mRNA each induced a significant increase in baseline mEPSC amplitude (sh-Ctrl vs. sh-Rai1 #1: n = 21-21, p=0.019, sh-Ctrl vs. sh-Rai1 #2: n = 18-19, p = 0.0011), without significantly altering mEPSC frequency or decay time (Fig 3A-D). Moreover, *Rai1*-KD induced a clear rightward shift in the cumulative probability distribution of mEPSC amplitudes in a manner that bears a striking similarity to changes in mEPSC distributions following chronic TTX treatment (Fig. 3E). An increase in surface expression of AMPA receptors (AMPARs) at synapses is a signature of synaptic upscaling following activity suppression. Consistent with previous observations, surface expression of the GluA1 AMPAR subunit at PSD-95-labeled excitatory synapses is significantly increased following chronic (24 hr) TTX treatment (sh-Ctrl Vehicle vs. TTX: n = 13-12, p = 0.0019, Fig. 3F). Likewise, we found a similar enhancement of surface GluA1 at synapses following 48 hr *Rai1*-KD (sh-Ctrl vs. sh-Rai1: n = 6-6 p = 0.0065, Fig 3F). Together, these results suggest that reduced *Rai1* expression induces functional changes in excitatory synaptic function that mimic synaptic upscaling induced by activity suppression.

**Figure 3.**
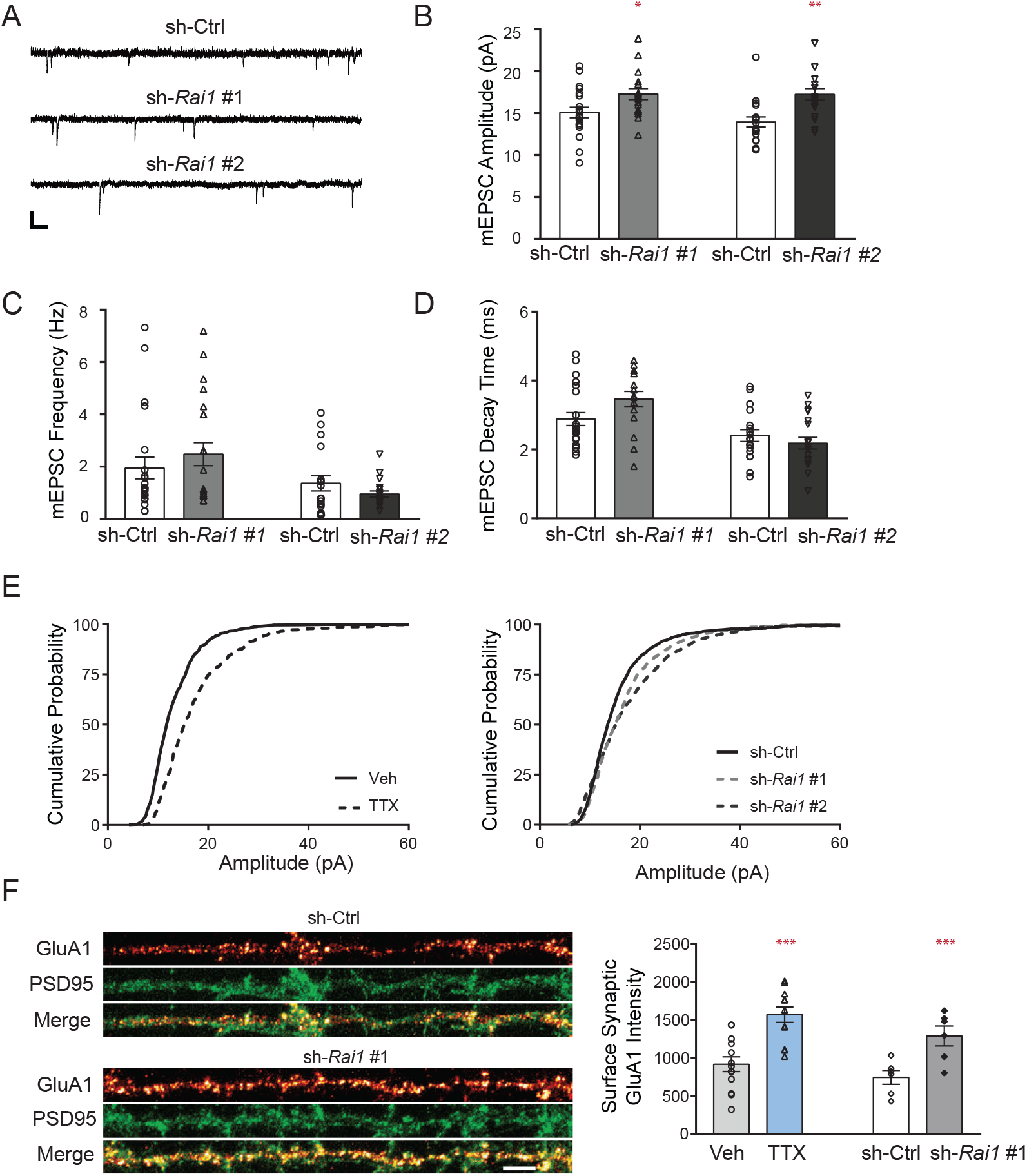
*Rai1*-KD increases the synaptic efficacy at baseline activity condition. **(A-D)** Example traces and mean ± SEM mEPSC amplitude (B), frequency (C), and decay time (D) for cultured rat hippocampal primary neurons recorded after transient transfection (48 hr) with either non-targeting shRNA (sh-Ctrl) or Rai1 targeting shRNA *(sh-Rai1* #1 or #2) at DIV12-14. Scale bar, 20 pA, 125 ms (sh-Ctrl, sh-Rai1 #1, n = 21-21, and sh-Ctrl, sh-Rai1 #2, n = 18-20) **(E)** Cumulative distribution of mEPSC amplitudes of sh-Ctrl transfected neurons treated by either vehicle or 1 μM TTX (left) and sh-Ctrl or sh-Rai1 transfected neurons (right). **(F)** Representative images of surface GluA1 (sGluA1, fire), PSD-95 (green) and sGluA1 & PSD-95 (merge) of sh-Ctrl and sh-Rai1 infected dendrites. Scale bar 10 μm. Bar graph of mean sGluA1 signal intensity in PSD-95 positive regions for vehicle or TTX (n = 13-12), and sh-Ctrl or sh-*Rai1* (n = 6-6) treated neurons. All bar graphs represent mean ± SEM, and comparisons between sh-Ctrl and sh-Rai1 were made with unpaired Student’s t-tests. *p < 0.05, **p < 0.01, ***p < 0.001.

### RAI1 promotes the transcriptional response to reduction in network activity

Having uncovered that RAI1 is essential to suppress the TTX-associated transcriptional program under baseline activity conditions (Fig. 2), we next tested if *Rai1*-KD has any impact on transcriptional induction and suppression upon TTX and BIC treatments. By calculating fold-changes of transcription, we found that *Rai1*-KD led to a significant impairment of transcriptional response to TTX, while transcriptional response to BIC was slightly weakened only for downregulation (Fig. 4A). However, in contrast to the 130 genes transcriptionally altered at baseline, DESeq2 gave only 8 genes as DE genes by *Rai1*-KD post TTX or BIC treatment, indicating that the impact of *Rai1*-KD is larger in resting neurons compared to drug-treated neurons (Fig. 4A, Table S3).

**Figure 4.**
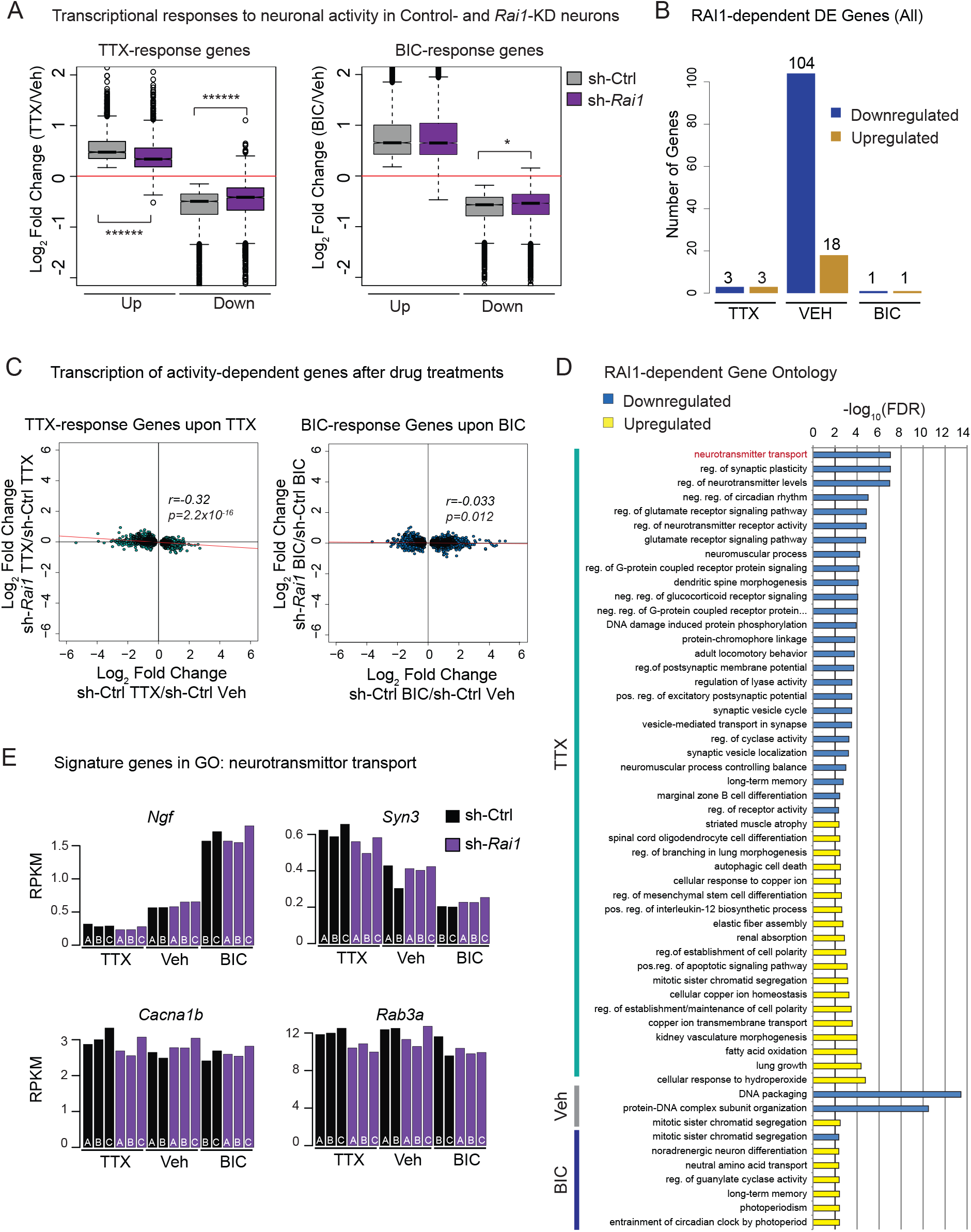
RAI1 positively regulates the transcriptional response to TTX. **(A)** The fold changes of TTX- and BIC-response genes in cultures treated by sh-Ctrl or sh-Rai1 (Wilcoxon rank-sum test). **(B)** Number of DESeq2-called differentially expressed genes (sh-Ctrl v sh-Rai1, p_adj_<0.05) after TTX, Vehicle, or BIC treatment. (D) RAI1-dependent gene ontologies (Biological Process, p_adj_ <0.005) discovered by RNA-Enrich (Lee et al., 2016) and filtered by ReviGO software (Supek et al., 2011). **(E)** RPKM values of four Sig-genes downregulated by *Rai1*-KD in the TTX-treated condition. The Sig-genes represent “neurotransmitter transport” (p_adj_ =9.3×10^-8^), the top-downregulated Biological Processes. The remaining Sig-genes are shown in Figure S7. Note slight but consistent inter-replicate changes upon *Rai1*-KD. r= Spearman’s rank correlation coefficient. In the box plots, whiskers represent 1.5 times IQR and the notch represents the 95% confidence interval of the median.

We then sought to determine if the strongly-impaired transcriptional response to TTX (Fig. 4A) was due entirely to the TTX-like transcriptional state of *Rai1*-KD culture at baseline or if RAI1 also contributes to the transcriptional response to TTX. Differential gene expression analysis by DESeq2 relies on an arbitrary statistical significance cutoff to report differentially expressed genes. However, the individual gene plot in the baseline condition revealed a global transcriptional trend resulting from small changes in many genes including those that failed to achieve statistical significance (Fig. 2E and S6B). To define the impact of RAI1 loss after TTX- and BIC-treatment, we therefore employed this individual-gene plot approach. We found that, after TTX treatment, the transcriptional changes of TTX-response genes in *Rai1*-KD cultures inversely correlate with their changes upon TTX treatment in the control condition (Fig. 4C, r=0.32, p=2.2×10^-16^, Spearman rank correlation coefficient, t-test). No correlation of TTX-response genes was observed after BIC treatment, suggesting RAI1 regulates TTX-associated transcription in baseline and TTX-treated conditions, but not under neural hyperactivation (Fig. S7A). Additionally, transcription of the BIC-response genes did not correlate with the transcription of genes in *Rai1*-KD cultures under any condition (Fig. 4A). Thus, *Rai1* -deficiency leads to subtle yet widespread impairment of the transcriptional response to TTX but not to BIC. Taken together, the Bru-seq results led us to conclude that 1) *RaI1*-deficiency shifts transcriptional profiles towards an activity-suppressed state in the resting network (Fig. 2), and 2) RAI1 is selectively required for the transcriptional response driven by network activity suppression (Fig. 3).

We sought to explore biological implications for such small but pervasive deficits in transcriptional response to TTX. We utilized RNA-Enrich, a gene ontology algorithm, in which the entire output of DESeq2 is analyzed, such that the program takes into account statistically-weaker changes in gene expression (Kim et al., 2012; Lee et al., 2016). Surprisingly, although *Rai1*-KD resulted in the greatest number of DE-genes in the vehicle-treated condition, RNA-Enrich identified many more RAI1-dependent biological processes after TTX treatment than vehicle-or BIC treatments (45 in TTX-, 3 in vehicle-, and 7 in BIC-treated cultures, Fig. 4B). The p_adj_ values were evidently lower in the post-TTX transcriptome data compared to BIC conditions (Fig. 4D). Furthermore, the RAI1-dependent gene ontologies after TTX treatment represent synapse-related processes, whereas those altered in the BIC and Vehicle-treated conditions show fewer ontologies directly relevant to neuronal activity (Fig. 4B and Table S4). The RNA-enrich provides the identity of signature genes, called Sig-genes, which significantly contributed to the enrichment of a given ontology (Kim et al., 2012; Lee et al., 2016). As expected, the genes that contributed to the enrichment of the synapse-related ontologies (e.g. *Ngf, Syn3, Cacna1b, Rab3a)* showed mild transcriptional changes upon *Rai1*-KD, yet the changes are consistent across biological replicates (Fig. 4E and Fig. S7B).

### Loss of RAI1 prevents synaptic upscaling but not downscaling

We next examined RAI1’s role in homeostatic synaptic scaling induced by chronic activity suppression (TTX, 24 hr) or chronic network hyperactivation (BIC, 24 hr). Consistent with the misregulation of TTX-responsive genes by *Rai1*-KD after TTX treatment (Fig. 4), we found that loss of RAI1 significantly impaired the induction of homeostatic upscaling during activity suppression in a cell-autonomous manner. Following transfection of scrambled or *Rai1* -targeted shRNAs (24 hr prior to TTX/BIC), we found that control neurons expressing the scrambled shRNA exhibited the normal increase in mEPSC amplitude 24 hr post-TTX (sh-Ctrl Veh vs TTX: n = 14-15, p = 0.0003, Fig. 6B) and the distribution of mEPSCs exhibited a clear and multiplicative rightward shift in cumulative frequency plots. By contrast, mEPSCs from neurons expressing either of two distinct *Rai1* shRNAs did not significantly increase following TTX exposure, and in fact, demonstrated a nominal trend for a decrease in amplitude (Fig. 5A-C). Despite this clear impairment of homeostatic upscaling, *Rai1*-KD had no effect on the induction of homeostatic downscaling following network hyperactivation with BIC (Fig. 5D-F). Both control neurons expressing scrambled shRNA and those neurons expressing *Rai1* shRNA exhibited significantly decreased mEPSC amplitudes (sh-Ctrl Veh vs BIC: n = 14-17, p = 0.0002, *sh-Rai1* #1 Veh vs. BIC: n = 9-8, p=0.026 and sh-Rai1 #2 Veh vs. BIC: n = 7-8, p = 0.0002, Fig. 6E), as well as a clear leftward multiplicative shift in mEPSC cumulative probability distributions. These results demonstrate that RAI is essential for homeostatic upscaling during activity suppression, but is otherwise dispensable for homeostatic downscaling during periods of network hyperactivation.

**Figure 5.**
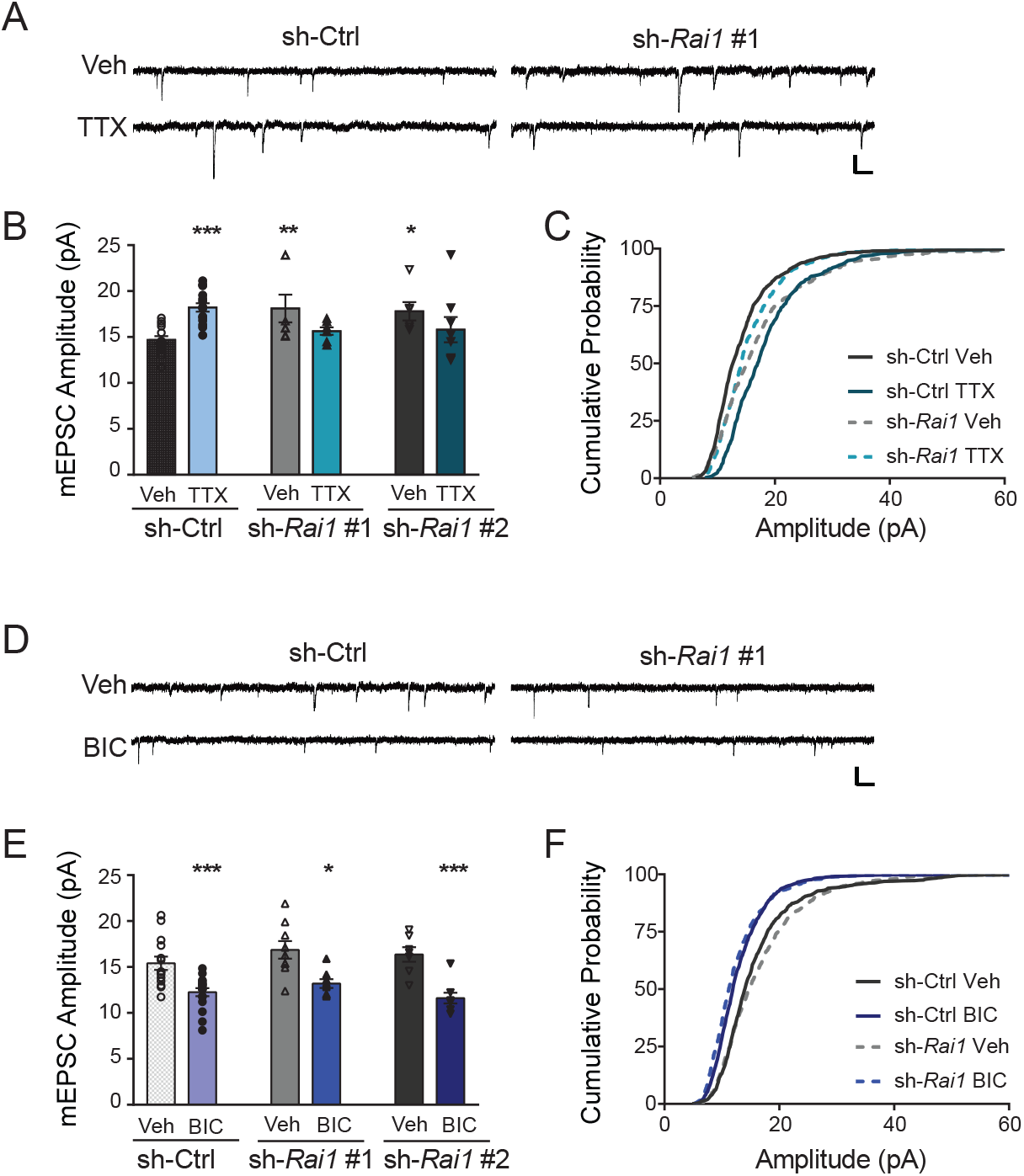
Rai1-KD impairs synaptic upscaling but not synaptic downscaling. **(A-C)** Representative mEPSC traces recorded from neurons transfected with either sh-Ctrl or sh-Rai1 and treated with either vehicle or 1 μM TTX. Scale bar, 20 pA, 125 ms. **(B)** mEPSC amplitude of sh-Ctrl, *sh-Rai1* #1 and *sh-Rai1* #2 treated either with vehicle or TTX (sh-Ctrl Veh, TTX n = 14-15, sh-Rai1 #1 Veh, TTX n = 7-7, sh-Rai1 #2 Veh, TTX n = 6-8). **(C)** Cumulative distribution of mEPSC amplitude of sh-Ctrl (black), *sh-Rai1* (teal; *sh-Rai1* #1 + #2) treated with vehicle (solid line) or TTX (dotted line). **(D-F)** Representative mEPSC traces recorded from neurons transfected with either sh-Ctrl or sh-Rai1 and treated with either vehicle or 10 μM BIC. Scale bar, 20 pA, 125 ms. **(E)** mEPSC amplitude of sh-Ctrl, *sh-Rai1* #1 and *sh-Rai1* #2 treated either with vehicle or BIC (sh-Ctrl Veh, BIC n = 14-17, sh-Rai1 #1 Veh, BIC n = 9-8, sh-Rai1 #2 Veh, BIC n = 7-8). (F) Cumulative distribution of mEPSC amplitude of sh-Ctrl (black), *sh-Rai1* (blue; sh-Rai1 #1 + #2) treated with vehicle (solid line) or BIC (dotted line). All bar graphs are represented as mean ± SEM. One-way ANOVA, followed by post-hoc Fisher’s LSD test were performed. *p < 0.05, **p < 0.01, ***p < 0.001.

**Figure 6.**
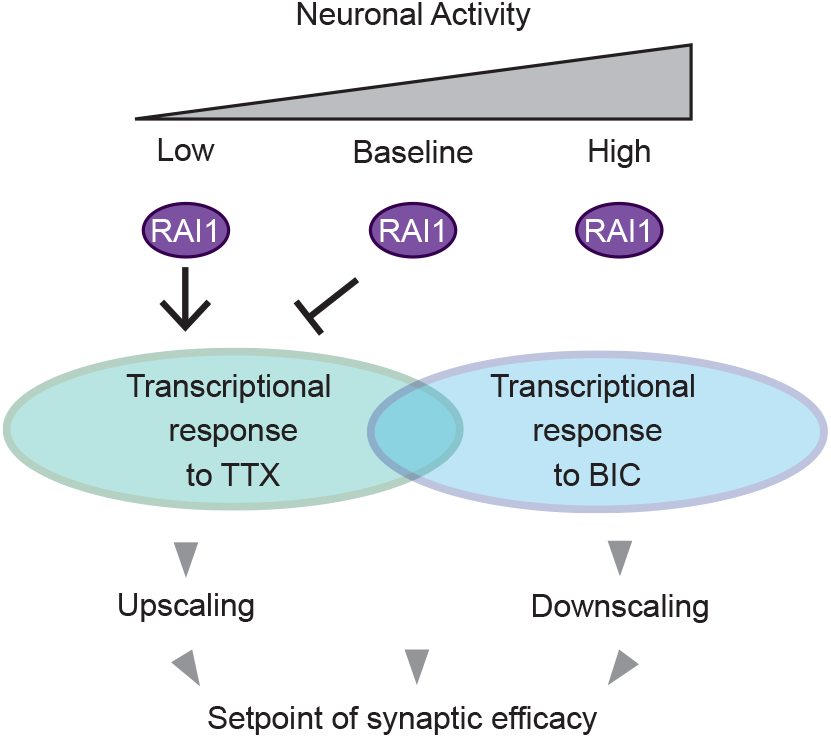
RAM’s role as a state-dependent transcriptional regulator of TTX-response genes: RAI1 alters synaptic efficacy through selective regulation of TTX-response genes under baseline and activity-suppressed states.

## Discussion

Our Bru-seq method and analyses have provided novel insights into activity-dependent transcription. We found widespread transcriptional responses to network activity shifts and its high sensitivity in detecting transcriptional downregulation. Furthermore, our data indicate that most dynamically regulated genes altered by hyperactivity or suppression are unique, not reciprocal. (Fig. 1). This is particularly interesting given that gene expression studies have tended to focus on bidirectional regulation of target genes (e.g. *Arc, Fos, Homer1, Bdnf)* (Okuno, 2011). Our results agree with a nascent proteome study on rat hippocampal neurons, in which the authors observed unique, common, and reciprocal changes in protein synthesis upon TTX and BIC treatments (Schanzenbächer et al., 2016). These observations suggest that in addition to the reciprocal transcriptional changes of key factors, distinct transcriptional mechanisms to low- and high-activity states may underlie upscaling and downscaling. Our data therefore provide a resource to begin exploration of distinct molecular machineries underlying homeostatic up- and down-scaling and regulation of bidirectional activity-dependent transcription.

Our data indicate that RAI1 is as a chromatin regulator that is selectively required for the transcription program of activity-suppression. Loss of RAI1 leads to misregulation of TTX-response genes, while leaving the uniquely BIC-genes unaffected (Fig. 2). In addition to its exclusive impact on TTX-response genes, an intriguing feature of RAI1 is its state-dependent roles in synaptic scaling. RAI1 deficiency shifts gene expression towards TTX-associated transcriptional states at baseline, and promotes the same transcriptional state once TTX is applied (Figs. 2 and 4). Under a hyperactivity condition, RAI1 is not required for transcription of TTX-associated genes (Fig. S7A, right panel). Since the correlation between *Rai1*-KD and TTX-associated transcription is stronger at baseline than TTX condition (baseline: r=0.53; TTX: r=0.32), RAI1’s influence on transcription appears greatest in the neurons without sensory inputs. In contrast to the RAI1’s roles in TTX-associated transcription, RAI1 is clearly dispensable for BIC-triggered transcription and synaptic downscaling at any activity state (Figs. 2 and 4). Thus, despite RAI1’s opposite roles in regulating TTX-associated genes between low- and baseline activity states, the ultimate role of RAI1 appears consistent — a specialized transcriptional regulator for TTX-associated genes (Fig. 6). This selective requirement of RAI1 is further corroborated with electrophysiological assessments (Figs. 3 and 5)

The mechanisms by which RAI1 performs this function remain to be resolved. RAI1 is a nucleosome-binding protein, which can act as a transcriptional coactivator (Bi et al., 2005; Burns et al., 2010; Carmona-Mora et al., 2012; Carmona-Mora et al., 2010; Darvekar et al., 2013; Elsea and Williams, 2011; Girirajan et al., 2009; Huang et al., 2016). Key genes for cognitive development, including *Bdnf*, are shown to be direct targets of RAI1 (Burns et al., 2010; Girirajan et al., 2009; Molina et al., 2008; Williams et al., 2012). Our Bru-seq data indicate that, under baseline levels of activity, RAI1 suppresses TTX-induced genes, while the same set of genes are positively regulated by RAI1 during activity suppression (Fig. 6). An integrative analysis of published ChIP-seq data suggest the majority of these genes are directly bound by RAI1 (Fig. S6A). It is plausible that RAI1’s role in transcription switches between a coactivator and a corepressor, in response to altered activity states. Alternatively, RAI1 binding to gene promoters might be dynamically regulated during activity shifts, as was the case with L3MBTL1 (Mao et al., 2018). TET3 and EHMT1/2 share their roles in homeostatic upscaling with RAI1 (Benevento et al., 2016; Yu et al., 2015). While TET3 positively regulates transcription by removing CpG methylation (Ito et al., 2010), EHMT 1/2 generally act as transcriptional repressors by placing the repressive histone mark, H3K9 methylation (Tachibana et al., 2005). How RAI1 functionally interacts with other chromatin regulators with roles in activity-dependent gene expression remains to be resolved.

It is noteworthy that the four previously-characterized chromatin regulators in synaptic scaling, i.e. Tet3, EHMT2, L3MBTL1 were all identified based on their expression changes or their target histone modifications by network activity shifts (Benevento et al., 2016; Mao et al., 2018; Yu et al., 2015). However, we found no indication that expression or sub-cellular localization of RAI1 is regulated by neuronal activity (Fig. S4–S5). The stable expression is similar to the case of EHMT1, in which EHMT1 levels remained constant while its binding partner EHMT2 levels varied in response to activity (Benevento et al., 2016). Chromatin regulators tend to show ubiquitous and stable expression. These observations raise a possibility that chromatin regulators that are implicated in human cognitive disorders could be involved in transcriptional response to activity shifts, even when their expression does not change during the process. Bru-seq and our analytical strategies employed in this study may prove useful to decipher how chromatin remodeling sculpts neural networks and plasticity.

How do RAI1’s new roles in synaptic scaling relate to cognitive function? As discussed earlier, RAI1 is implicated Smith-Magenis Syndrome (SMS) (Bi et al., 2004; Girirajan et al., 2005; Slager et al., 2003) and mouse models of heterozygous and homozygous *Rai1*-KO displayed learning deficits, abnormal circadian behavior, altered social behavior, and obesity (Bi et al., 2005; Bi et al., 2007; Huang et al., 2018; Lacaria et al., 2013). Thorough characterization of cell-type specific *Rai1*-KO mice attributed the learning deficits to GABAergic interneurons rather than glutamatergic excitatory neurons (Huang et al., 2016). Since our mEPSC recording was performed in the pyramidal cells that incorporated *Rai1*-shRNA plasmid, and the transfection efficiency is low (~0.1%), RAI1’s role in synaptic scaling is likely cell-autonomous to excitatory neurons. The lack of learning deficits in forebrain-specific *Rai1*-KO (Emx1-Cre: Rai1^flox/flox^) may suggest that RAI1’s roles in synaptic scaling is irrelevant to cognitive function. Alternatively, synaptic scaling deficits by acute *Rai1* depletion might be compensated during development by unknown genes. The mice lacking RAI1 only in excitatory neurons may undergo compensatory neurodevelopment processes that involve RAI1-positive cell types, in which case the mouse model may not reflect neurodevelopmental deficits in human that is caused by *RAI1* heterozygosity in all cells. Additionally, the recent finding of reduced dendritic spine density in the prefrontal cortex of 4 week old *Rai1*-heterozygous mice, (Huang et al., 2018) may be result in part from impaired upscaling in RAI1-deficient excitatory neurons during development.

In addition to cognitive function, sleep-wake cycle is another behavior in which RAI1’s roles in synaptic scaling might be involved. Sleeping problems of SMS patients have been associated with inverted circadian rhythms (Boone et al., 2011; Elsea and Williams, 2011; Gropman et al., 2006; Potocki et al., 2000; Williams et al., 2012). Regulation of *CLOCK* gene by RAI1 has provided a molecular mechanism, which potentially explains the inverted circadian rhythm (Williams et al., 2012). Meanwhile, accumulating evidence has indicated that the brain undergoes synaptic downscaling during sleep, which normalizes the strengthened synaptic connection by experience during the awake state (Kuhn et al., 2016) (de Vivo et al., 2017; Diering et al., 2017). Our work demonstrates that reduced network activity elicits unique transcriptional responses (Fig. 1). Together, these observations predict that, during sleep, network activity becomes low due to fewer sensory inputs, reminiscent of TTX-treated neurons, which would trigger transcriptional responses that in turn upscale synapses during the awake state. It is tempting to speculate that RAI1-dependent synaptic upscaling might contribute to the higher synaptic efficacy during the awake state. Future studies should address how activity-dependent transcription and circadian gene oscillation interact, where RAI1 acts, and how disruption of these processes underlies cognitive and/or sleeping issues of SMS and related cognitive deficits.

In addition to Bru-seq, several nascent RNA-seq approaches have been employed to examine gene expression in neurons. Schaukowitch *et al.* adopted Gro-seq, a nuclear run-on assay coupled with deep sequencing, to profile transcriptional activity in TTX-treated neurons (Schaukowitch et al., 2017). Gro-seq involves isolation of nuclei and incorporation of BrUTP during the run-on reaction *in vitro* (Core et al., 2008). We found that Gro-seq is more advantageous to detect unstable RNAs, such as eRNAs, compared to Bru-seq (Agarwal et al., 2017). The highly-sensitive detection of unstable RNAs is likely due the lack of active exosomes in the run-on reaction, which however points to a potential risk to observe *in vitro* artifacts. In contrast, Bru-seq faithfully monitors transcription within the cells at a given moment (Paulsen et al., 2014; Paulsen et al., 2013). A limitation of the Bru-seq approach is the lack of cell-type specificity. Recently, Zajaczkowski *et al.* reported nascent RNA-sequencing specifically in neurons that were depolarized by KCl (Zajaczkowski et al., 2018). In this approach, neuron-specific RNA labeling was achieved by the *Synapsin I* promoter-driven expression of a *Toxoplasma gondii* enzyme, uracil phophoribosyltransferase (UPRT). UPRT enables incorporation 5-ethynyl-uracil (5EU) into RNA. The labeled RNAs were biotinylated, isolated, and subjected for sequencing (Cleary et al., 2005). The UPRT-5EU system identified over 3,000 depolarization-regulated genes over the 3 hr KCl treatments, which likely detected both nascent transcripts and steady-state mRNAs. Cell-type specificity of the UPRT-5EU system comes with the cost of introducing the UPRT transgene and additional experimental steps to label RNAs. Recent transcriptome studies of brain cell types and single-cell RNA-seq have allowed us to retrospectively attribute the transcriptional changes in Bru-seq data to certain cell types to some extent (Fig. S2). Thus, one can choose the most suitable experimental approach for nascent RNA profiling depending on their goals of the study.

## Supplementary Figure legends

**Figure S1.**
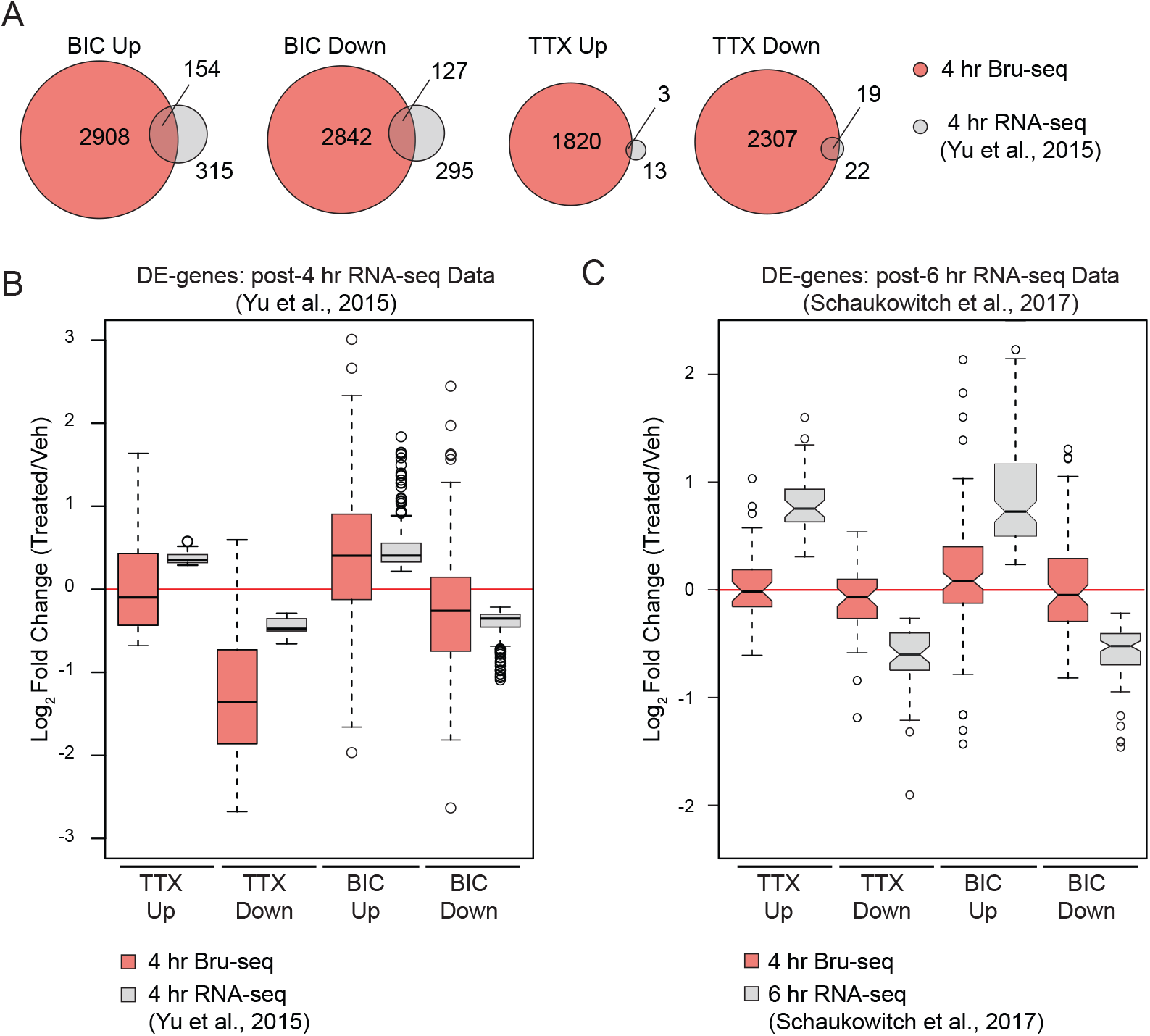
Bru-seq vs. mRNA-seq. **(A)** Overlap of DE genes in Bru-seq and mRNA-seq. DE genes were called using identical DESeq2 parameters. mRNA-seq datasets were obtained from from (Yu et al., 2015). **(B)** Expression changes of DE-genes upon BIC and TTX treatments in Bru-seq and mRNA-seq (4 hr post-treatment) (Yu et al., 2015). **(C)** Comparison of Bru-seq data and mRNA-seq data of 6 hr post-treatment (Schaukowitch et al., 2017).

**Figure S2.**
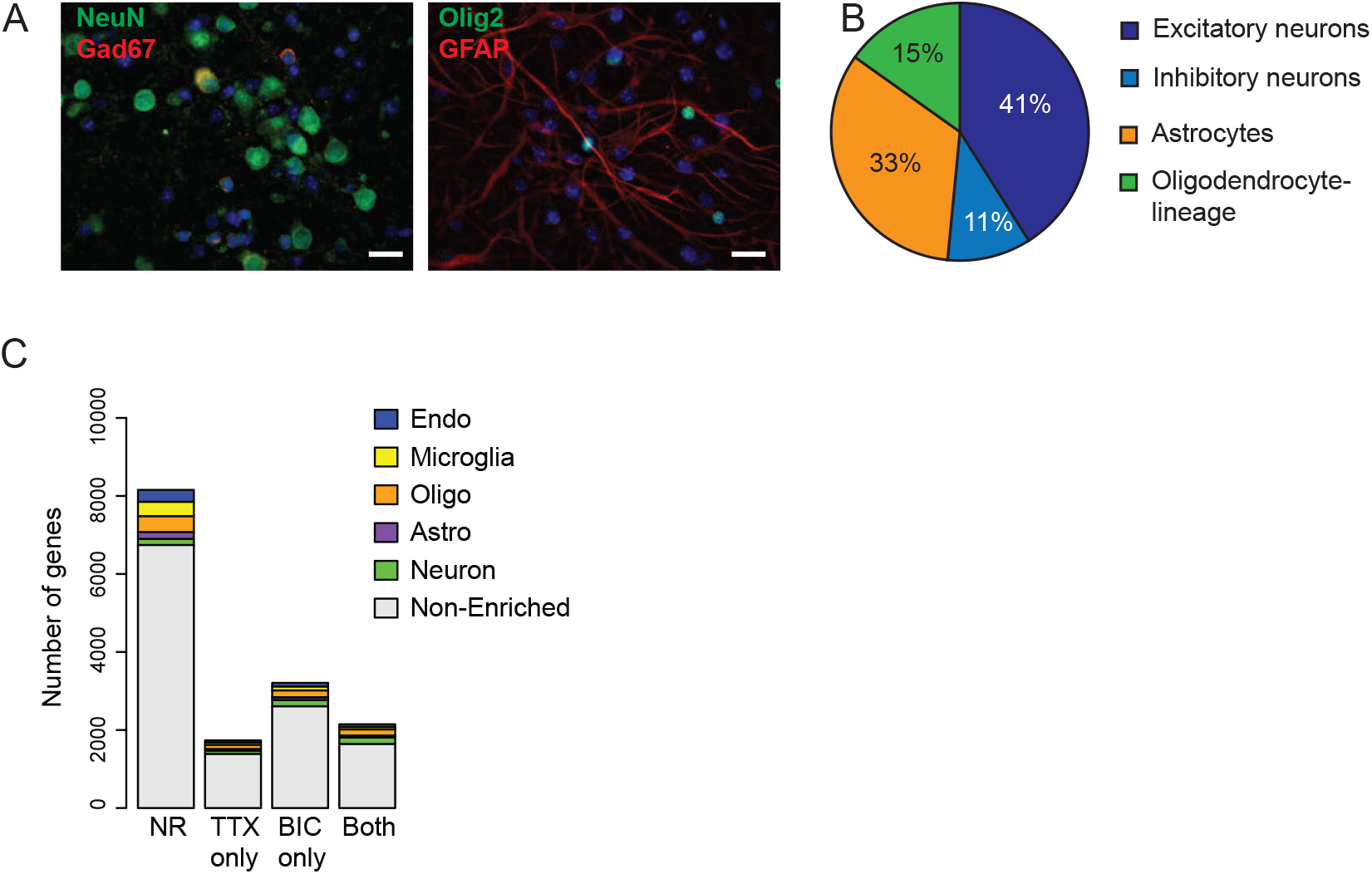
Cell-type analysis of the primary forebrain neuron cultures. **(A)** Representative immunofluorescence images of the primary forebrain neuron culture (DIV17) with antibodies against NeuN, Gad67, GFAP, or Olig2. Nuclei were visualized by DAPI. Scale bar: 20 μm. **(B)** Quantification of cell types. Cell types were determined as follow. Excitatory neurons; NeuN(+), Gad67(-), Inhibitory neurons; NeuN(+), Gad67(+), Astrocytes; NeuN(-),GFAP(+), Olig2(-). 33.5%, and Oligodendrocyte lineage cells; NeuN(-), Olig2(+). We did not observe cells with CD11b, a microglia marker. Each cell type was calculated as the % of all DAPI+ cells and shown as an average of two biological replicates. **(C)** The number of cell type-specific genes and their response to TTX or BIC in the Bru-seq data. Cell type-specific genes were obtained from mRNA-seq data of separated cells by immunopanning of P7-P17 mouse cortices (Zhang et al., 2014). Oligo: cells within the oligodendrocyte lineage. NR: Non-responsive genes. Both: Genes that respond to both TTX and BIC.

**Figure S3.**
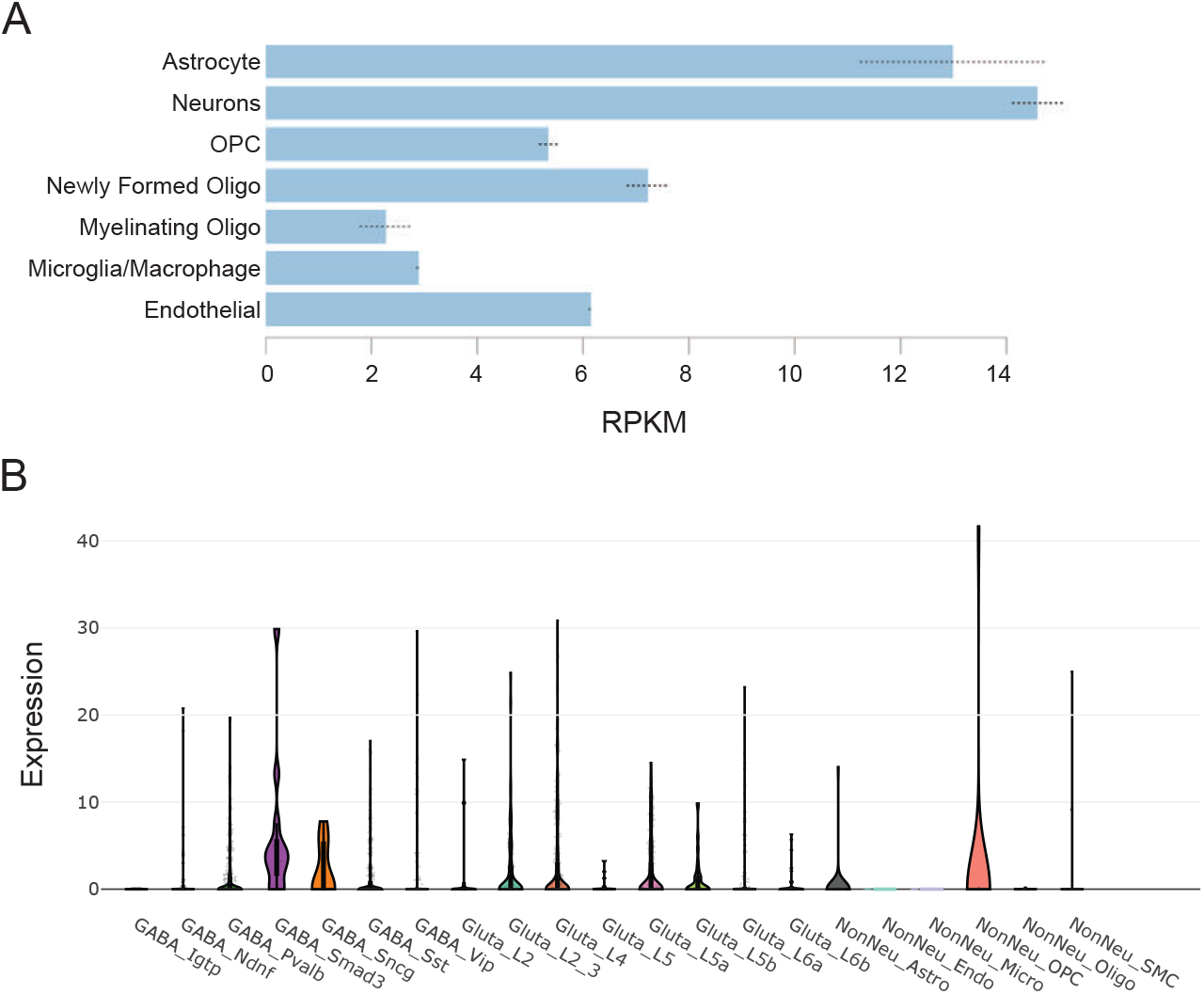
RAI1 expression in publicly-available datasets. **(A)** *Rai1* mRNA levels in neurons and non-neuronal cells of the adult mouse cortices (Zhang et al., 2014). **(B)** *Rai1* mRNA levels in single cell mRNA-seq data of mouse visual cortex (Tasic et al., 2016).

**Figure S4.**
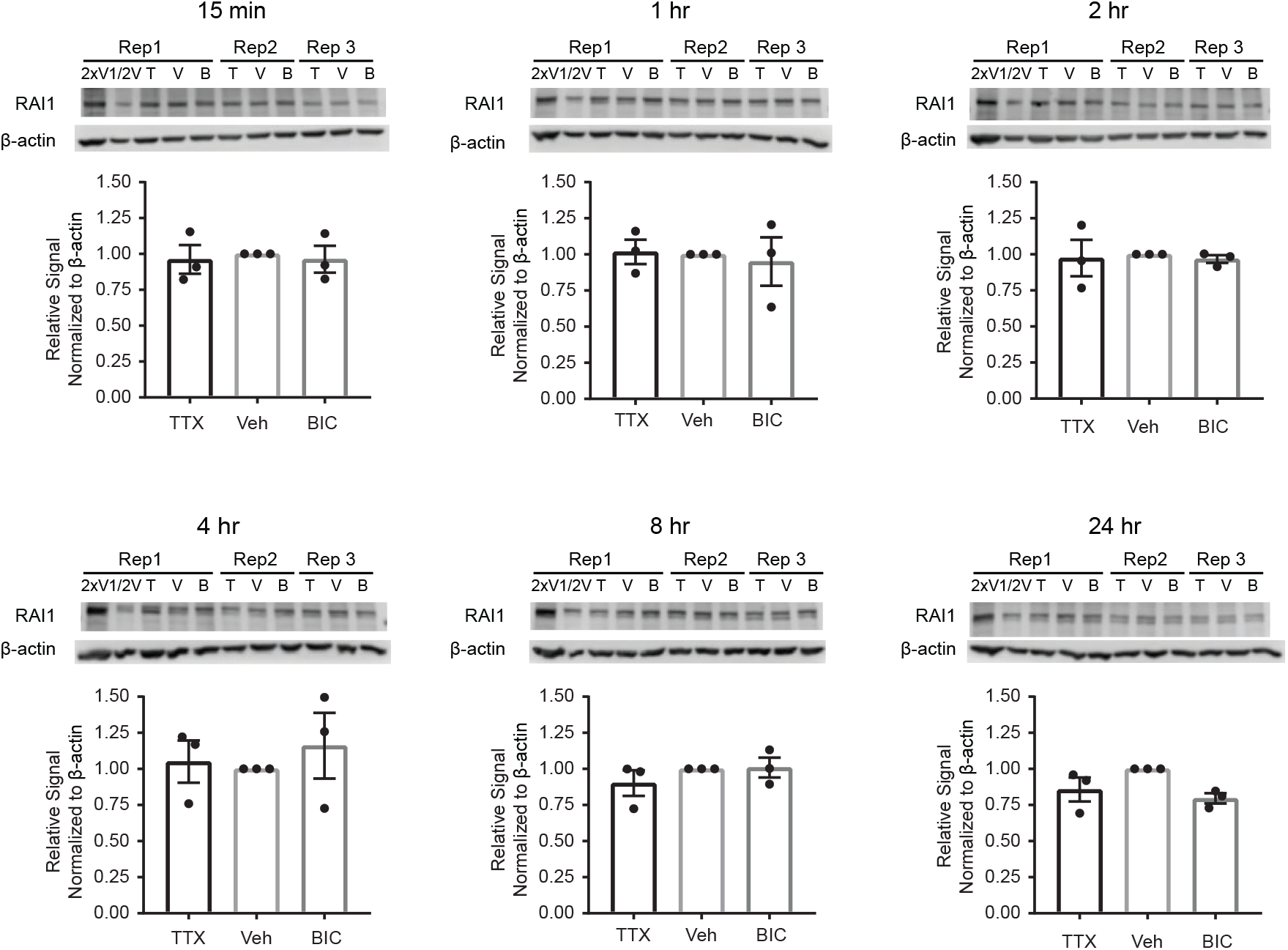
RAI1 protein level during neuronal-activity shifts. Rat cortical neurons (DIV14) were treated with TTX (T), BIC (B) or vehicle (V) for the indicated times. Cells were harvested in 3biological replicates and analyzed by Western blot with a custom anti-RAI1 antibody. The 270-kDa full-length RAI1 protein band intensity was visualized and quantified in the linear range using LI-COR C-Digit and Image Studio software. Statistical significance was evaluated with one-way ANOVA. No treatment reached p<0.05.

**Figure S5.**
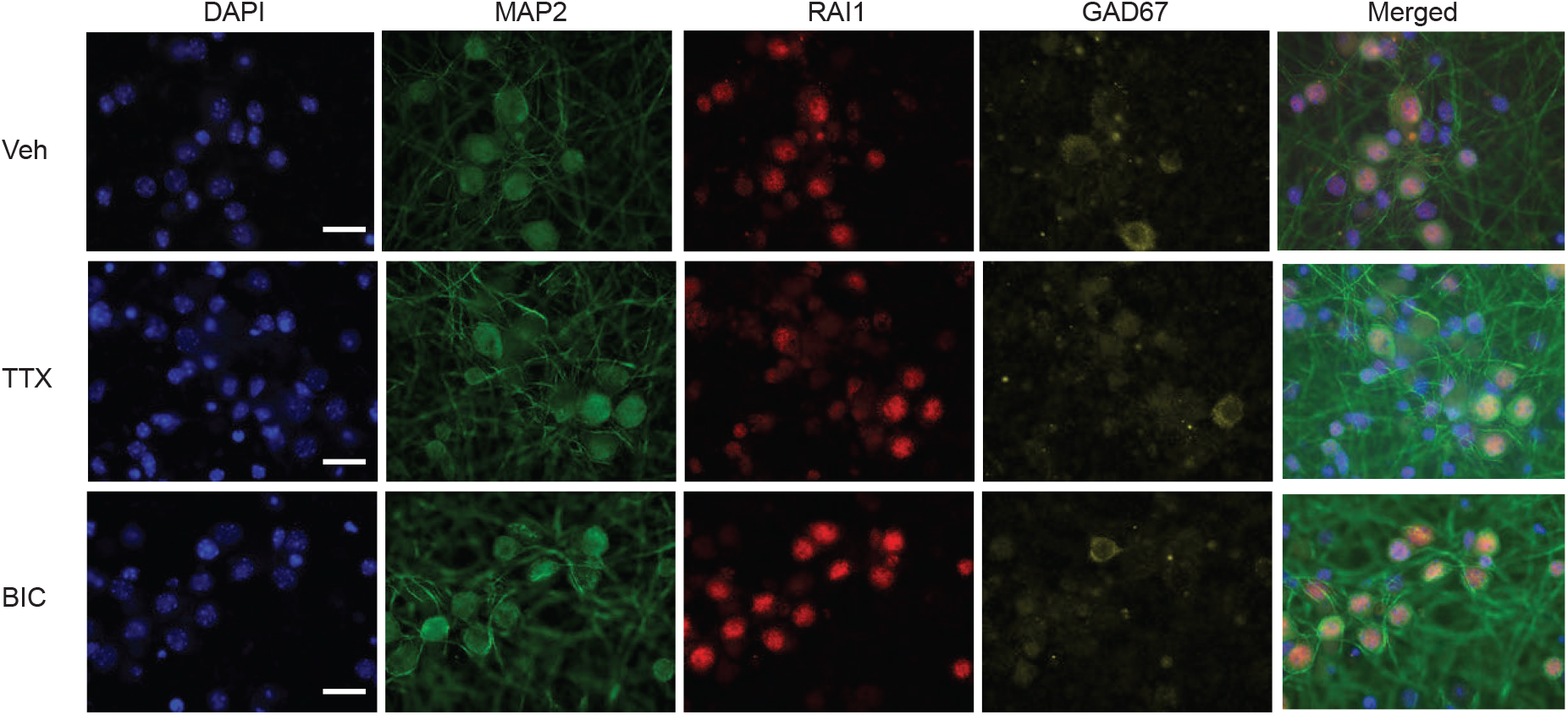
Sub-cellular localization of RAI1 during neuronal-activity shifts. Sub-cellular RAI1 localization was assessed by immunofluorescence in the mouse forebrain neuron culture (DIV17) using an anti-RAI1 antibody. RAI1 displayed nuclear localization in excitatory and inhibitory neurons. RAI1 did not show any sub-cellular or sub-nuclear localization by TTX and BIC treatments for 4 hr. We obtained similar results at other timepoints (15 minutes, 1 hr, 2 hr, 8 hr, and 24 hr, data not shown).

**Figure S6.**
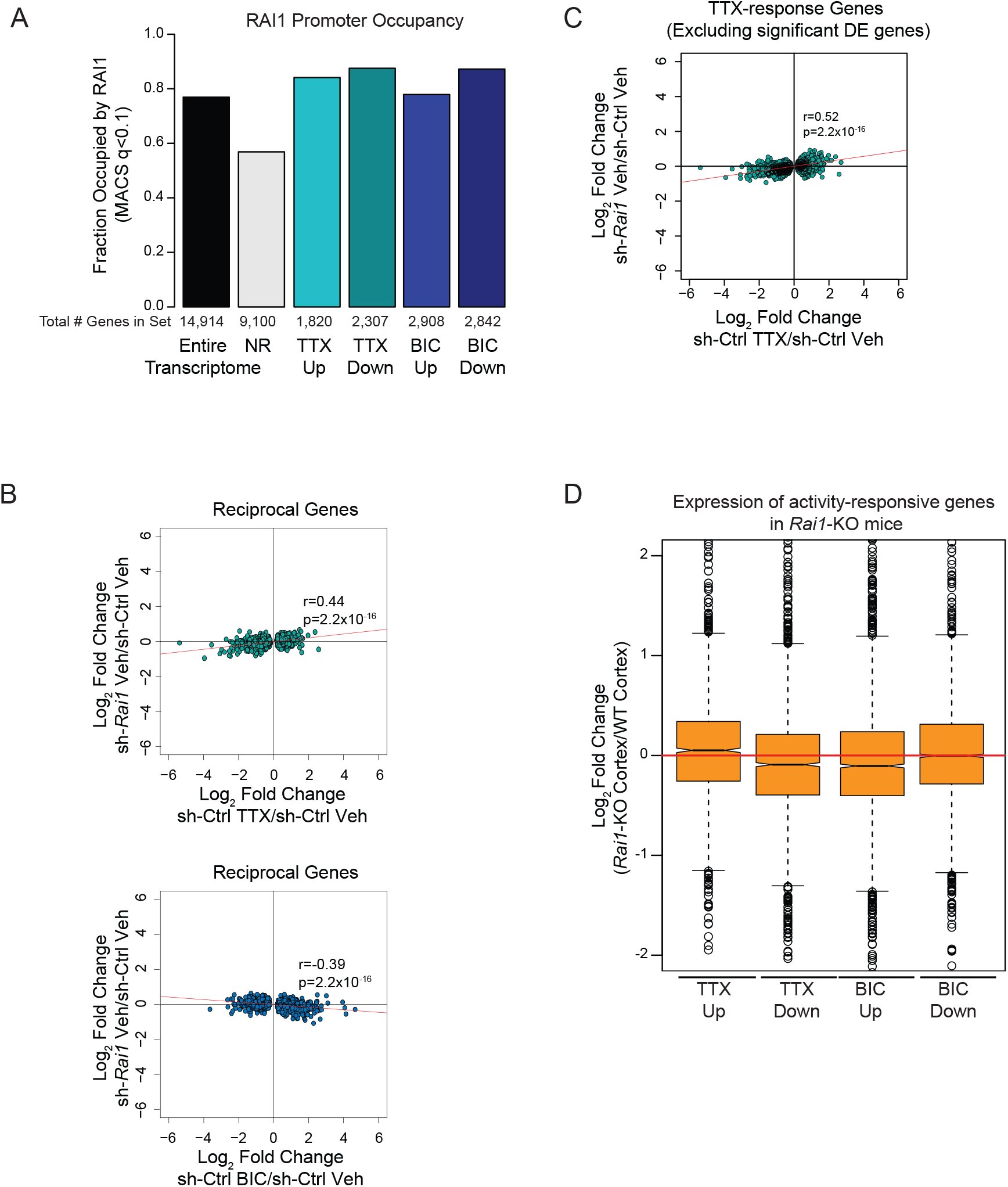
Integrative analysis of the Bru-seq, RAI1 ChIP-seq, and mRNA-seq data of *Rail-knockout* mice. **(A)** RAI1 occupancy at the promoters of TTX- and BIC-response genes. RAI1 ChIP-seq data were obtained from the cortices of 8-week old mice (Huang et al., 2016). RAI1 occupies ~80% of activity-dependent genes with no apparent enrichment in any group. Promoters were defined by ± 1 kb of annotated transcription start sites and overlap with RAI1 ChIP-seq peaks were computed using MACS2 (Zhang et al., 2008). **(B)** Transcription of reciprocal genes in the *Rai1*-KD culture at baseline show a positive correlation with TTX-treated transcriptome and a negative correlation with BIC-treated transcriptome of the normal culture. **(C)** Premature TTX-response of *Rai1*-KD culture is still observed after excluding the *Ral1*-dependent genes at baseline. **(D)** Expression of TTX- and BIC-response genes in the panneuronal *Rai1*-knockout cortex. mRNA-seq data were obtained from 3 week-old Rai1^flox/flox^: *Nestin-Cre* and control mice (Huang et al., 2016). Fold changes in KO vs Control mice were calculated using the RPKM values. The *Rai1*-KO mRNA expression shows a trend of TTX-treated transcription states.

**Figure S7.**
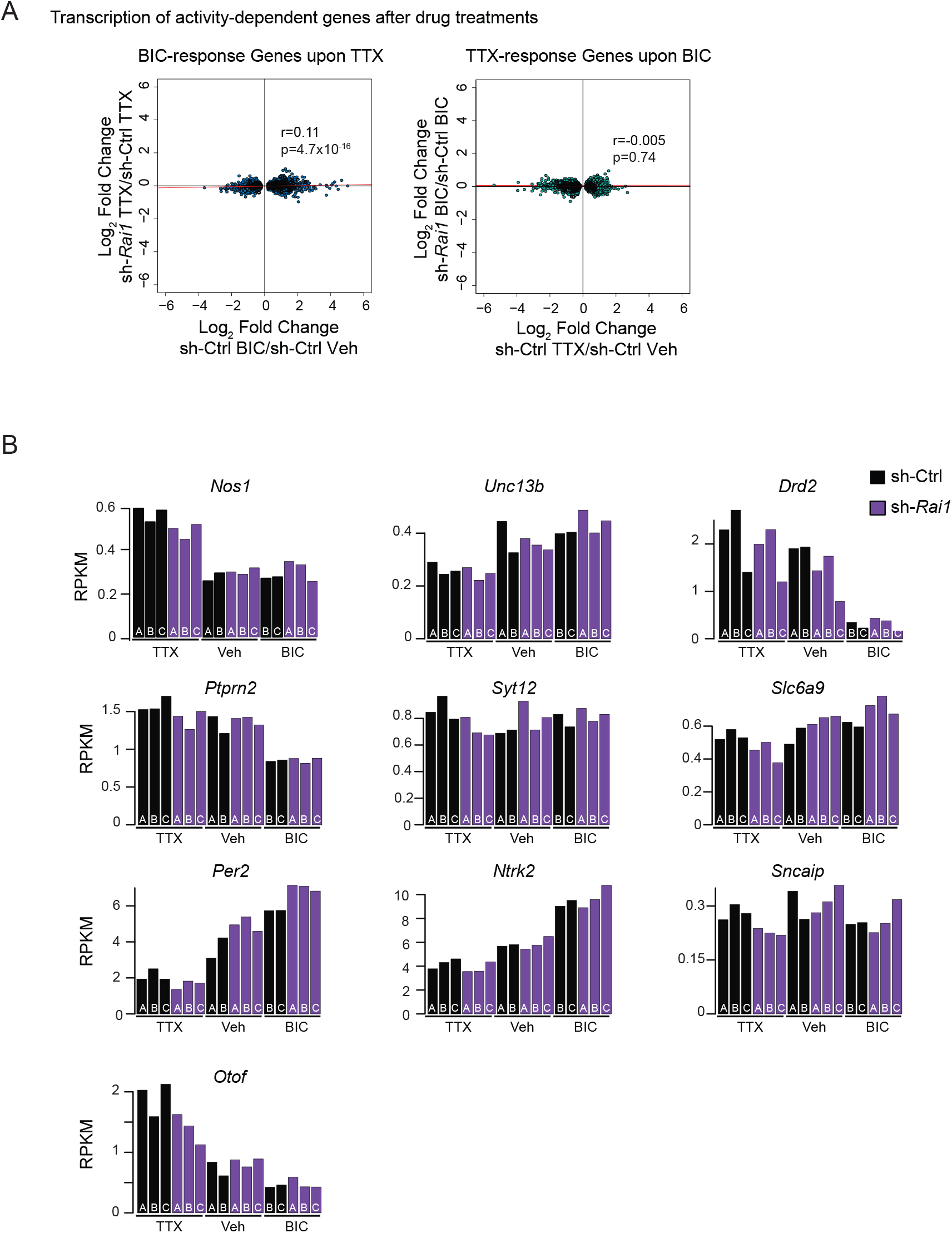
Impact of RAI1 loss on transcription after TTX- or BIC-treatments. **(A)** The fold changes of TTX- and BIC-response genes by *Rai1-KD* after BIC or TTX treatment. **(B)** RPKM values of Bru-seq data are represented for Sig-genes of “neurotransmitter transport”, which was downregulated by *Rai1*-KD in the post-TTX condition (p_adj_: 9.3 x 10^-8^). Note consistent reduction in RPKM across biological replicates in *Rai1*-KD culture in the TTX-treated condition.

## Materials and Methods

### Primary neuron culture and shRNA-mediated *Rai1*-KD

The cortices and hippocampi from E18.5 mouse pups were pooled into biological replicates with identical female-male ratios. Sex of the pups was determined by PCR using primers for the *ZFY* gene (Table S5). Primary culture of neurons was carried out as previously described (Iwase et al., 2016) (Vallianatos et al., 2018). Briefly, dissociated tissues were plated at 4 million cells/6 cm poly-D-lysine-coated plate (Sigma) grown in Neurobasal Media supplemented with B27 (Gibco, #17504044). No mitotic inhibitors were added, allowing the growth of non-neuronal cells. Half the culture medium was freshened every 3-5 days. On DIV 14, cells were infected with lentiviral shRNA as previously described (Vallianatos et al., 2018). Lentivirus were generated using co-transfection into HEK-293t cells of psPAX2 (Addgene, 12260), pMD2.G (AddGene, 12259) and pLKO plasmids containing shRNA against *Rai1* untranslated region (Rai1-shRNA #1: Sigma, TRCN0000124984) or coding region (Rai1-shRNA #2: Sigma, TRCN0000328334) or scramble shRNA (Sigma, SHC202). For Bru-seq experiments, we used SHC202 and *Rai1*-shRNA #1. For electrophysiology, we used SHC202, *Rai1*-shRNA #1 and #2, whose target sequences are identical between mouse and rat. The conditioned media containing lentiviruses was collected, concentrated with Lenti-X concentrator (Takara, 631232), and resuspended in Neurobasal medium, and stored at −80°C. The titer of lentivirus was determined by survival of transduced 293 cells under puromycin and a comparable amount of virus that result in >90% survival of infected neurons was used for all biological replicates. Puromycin was not added to cultured neurons for experiments.

### Network activity alterations and Bru-seq experiments

On DIV17, cells were treated with bicuculline-methiodide (Abcam, ab120108, 20 μM), TTX (Tocris, 1069, 1 μM), or vehicle (sterile water), for 4 hours. 3 hours and 40 minutes post treatment, bromouridine (Bru, Sigma, dissolved in PBS) was added to cultures at 2 mM final concentration. Cultures were harvested in Tri-reagent BD (Sigma, T3809) and frozen immediately. RNA was purified using phenol-chloroform extraction and isopropaonol precipitation, treated with DNAse-I (NEB) then fragmented by high-magnesium, high temperature incubation. From 1 μg of total RNA, enrichment of Bru-containing RNA and library preparation were performed as previously described (Paulsen et al., 2014; Paulsen et al., 2013) with minor modifications. We designed custom adaptors (Table S5) which were directly ligated to the 3’ ends of RNA using RNA ligase 1 (NEB Cat. No. M0437) and truncated RNA ligase KQ (NEB M0373). Bromouridine-labeled RNAs were immunoprecipitated using anti-BrdU antibody (Santa Cruz Biotechnology, sc-32323). Enriched RNAs were reverse transcribed using a primer complementary to the RNA adaptor (Table S5). Adaptor duplexes with 5- or 6-base pair random nucleotide overhangs were ligated to the 3’ end of the cDNA (Table S5). The cDNA libraries were amplified using primers that carry Illumina indices, then 180-400 bp DNA fragments were isolated using by an agarose gel. The nucleotide sequences of primers used for library amplification are found in Table S5. The libraries were subjected to single-end 50-bp sequencing using IIlumina HiSeq 2000 platform. We performed 2 to 3 biological replicates for all drug treatment and knockdown conditions.

### Sequencing data analysis

After confirming the quality of sequencing data by FastQC, reads were mapped to mm9 reference genome using Bowtie2 (Langmead and Salzberg, 2012) and annotated with Tophat2 (Kim et al., 2013). Adaptors were trimmed using BBDUK (http://jgi.doe.gov/data-and-tools/bb-tools/), when 2-30 bp on the left of the read matched the predicted adaptor (k=30, mink=2, minlength=15, hdist=1). Bru-seq signals were quantified by FeatureCounts (Liao et al., 2014). We excluded *Rn45s, Lars2, Rn4.5s, Cdk8, Zc3h7a* and the mitochondrial chromosome to avoid counts of overamplified genes that may skew RPKM normalization. DE-genes were identified using DESeq2 (Love et al., 2014) using the same parameters for the Bru-seq data and three published mRNA-seq datasets of neuron culture and *Rai1*-KO mice (Huang et al., 2016; Schaukowitch et al., 2017; Yu et al., 2015). We also used DESeq2 to calculate RPKM expression values across the entire genic regions, including introns. Gene ontology was examined using RNA-Enrich (Lee et al., 2016). Significance cutoff for reporting Sig-genes was an unadjusted p value< 0.05. We only presented GO terms that contain 5 to 250 genes.

### Western blot

To validate *Rai1*-KD in mouse forebrain neuron culture, *Rai1*-KD and control cultures were harvested at 3 days after lentiviral transduction and subjected to Western blot analysis as described previously (Iwase et al., 2016). RAI1 antibodies were generated by immunizing rabbits with a synthesized RAI1 peptide (aa 28 to 42, ENYRQPGQAGLSCDR, Thermo Fisher Scientific), followed by affinity purification using the peptide as the affinity ligand (Thermo Fisher Scientific). Anti-PCNA antibody (Santa Cruz sc-56, 1:1000) was used for a loading control. For analysis of RAI1 level during activity shifts, the cortices from E18.5 rat pups were dissected, dissociated, and plated at 700,000 cells/well in a PDL-coated 6-well dish. Neurons were grown in Neurobasal/B27 medium for 14 DIV. Vehicle (1% water), TTX (1 μM) or BIC (20 μM) were added to the culture and cells were harvested with a 1:1 mixture of 2X Laemmeli buffer (BioRad, 1610737, 1:20 beta-mercaptoethanol) and radioimmunoprecipitation assay (RIPA) buffer supplemented with 50 mM BGP and 1 mM Na3VO4. Protein samples were boiled for 10 minutes at 100°C. 10-15 μg of each sample was loaded per lane, separated by 7.5% SDS-PAGE, and transferred onto PVDF membrane (Millipore IPVH00010). Membranes were then blocked with 5% skim milk or 3% blotting-grade blocker (BioRad 1706404) for 1 hr, probed overnight with the following primary antibodies diluted in 3% BSA (Fisher Scientific BP1600): RAI1 (1:1000), beta-actin (Sigma A5441, 1:20,000). Horseradish peroxide (HRP)-conjugated secondary IgG antibodies (EMD Millipore AP132P or AQ160P) were also diluted in 3% BSA, and the HRP signal was developed with various chemiluminescent substrates from Thermo Fisher Scientific (34080 or 34095) and Li-COR Biosciences (926-95000). Protein band intensity was visualized and quantified in the linear range using LI-COR C-Digit and Image Studio software. Results were compared using one-way ANOVA.

### Immunocytochemistry

Two biological replicates of forebrain neuron cultures were obtained from E17.5 mouse embryos. On DIV19, they were fixed with 4% paraformaldehyde in 16% sucrose/PBS, permeabilized with 0.25% Triton-X in 1X PBS, blocked for 30 minutes with 10% bovine albumin serum (Sigma A2153), and overnight incubation of antibodies in 3% BSA at 4°C. Primary antibodies used in the study are following. anti-NeuN (EMD Millipore, MAB377, 1:1000), anti-GFAP (NeuroMab N206A/8, 1:1000), anti-MAP2 (EMD Millipore, AB5543, 1:1000), anti-OLIG2 (EMD Millipore, AB9610, 1:1000), anti-CD11b (Abcam, ab133357, 1:500), anti-GAD67 (Santa Cruz, sc-5602, 1:1000). Secondary antibodies (Invitrogen Alexa Fluor 488, 555, or 647) were applied for 45 min at room temperature. Fluorescence images were acquired using an Olympus BX61 fluorescence microscope (60X oil-immersion lens) and CellSense software. Immunoreactivity was quantified semi-automatedly using a custom ImageJ script after confirming specific staining by visual inspection.

### Electrophysiology

All animal use followed NIH guidelines and was in compliance with the University of Michigan Committee on Use and Care of Animals. Dissociated postnatal (P0-2) rat hippocampal neuron cultures were prepared as previously described (Henry et al., 2012). Neurons were transfected with 1.0 μg of scrambled or *Rai1*-shRNA-expressing plasmids with the CalPhos Transfection kit (ClonTech) or Lipofectamine 2000 (ThermoFisher Scientific) according to the manufacturer’s protocols. All experiments were performed 48 hours after transfection. mEPSCs were recorded from a holding potential of – 70 mV with an Axopatch 200B amplifier from neurons bathed in HEPES buffered saline (HBS) containing: 119 mM NaCl, 5 mM KCl, 2 mM CaCl2, 2 mM MgCl2, 30 mM Glucose, 10 mM HEPES (pH 7.4) plus 1 μM TTX and 10 μM bicuculine; mEPSCs were analyzed with Synaptosoft MiniAnalysis software. Whole-cell pipette internal solutions contained: 100 mM cesium gluconate, 0.2 mM EGTA, 5 mM MgCl2, 2 mM ATP, 0.3 mM GTP, 40 mM HEPES (pH 7.2). Statistical differences between experimental conditions were determined by unpaired Student’s t-tests (Fig. 3) or one-way ANOVA followed by post-hoc Fisher’s LSD test (Fig. 6).

### Surface GluA1 expression analysis

Surface GluA1 staining was conducted as previously described with slight modification (Henry et al., 2012). On DIV12, rat cultured hippocampal cells were infected either with lentivirus carrying sh-Ctrl or sh-Rai1 as described above. After 48 hours of incubation, cultured cells were live-labeled with rabbit anti-GluA1 antibody (EMD Millipore, ABN241, 1:1000) for 20 min at 37°C, fixed with 2% paraformaldehyde, and further labeled with mouse anti-PSD-95 (EMD Millipore, MAB1596, 1:1000). Goat anti-mouse Alexa 488 and Goat anti-rabbit Alexa 555 secondary antibodies (Abcam, 1:500) were applied for 60 min at room temperature to visualize PSD-95 and GluA1 staining. Images of PSD-95 and GluA1 were aquired using an inverted Olympus FV1000 laser-scanning confocal microscope using a Plan-Apochromat 63 X /1.4 oil objective. Then, synaptic GluA1 was defined as a particle that occupied > 10% of the PSD-95 positive area, and the average integrated intensity of synapatic GluA1 was calculated using a custom macro for ImageJ. Statistical differences between experimental conditions were determined by unpaired Student’s t-tests.

### Data Availability

Sequencing data generated for this study have been submitted to the NCBI Gene Expression Omnibus (GEO; http://www.ncbi.nlm.nih.gov/geo/) under accession number GSE121749.

## Supporting information

Supplemental File 1_TTX

Supplemental File 1_BIC

Supplemental File 2_TTX

Supplemental File 2_BIC

Supplemental File 3_Vehicle

Supplemental File 3_TTX

Supplemental File 3_BIC

Supplemental File 4_Vehicle

Supplemental File 4_TTX

Supplemental File 4_BIC

## Acknowledgements

We thank Dr. Saurabh Agarwal for providing technical support for Bru-seq experiments and scripts for mapping and annotating RNA-seq data. P.M.G. was supported by an NSF Graduate Research Fellowship Program (DGE #1256260). T.T. was supported by PRISMS post-doctoral fellowship. A.C. was supported by an NIH NRSA fellowship (18-PAF03228). M.A.W. was supported by a Summer Fellowship from the University of Michigan Language, Sciences and Arts Honors Program. This work was supported by the University of Michigan Medical School (to S.I.), March of Dimes Foundation (to S.I.), NIH (R01NS089896 and R21NS104774 to S.I.; R01NS097498 to M.A.S), the Farrehi research fund (to S.I.). The authors thank the members of Iwase and Sutton laboratories for discussion and support.

## References

Abbott, L.F., and Nelson, S.B. (2000). Synaptic plasticity: taming the beast. Nat Neurosci 3, 1178.

Agarwal, S., Garay, P.M., Porter, R.S., Brookes, E., Murata-Nakamura, Y., Macfarlan, T.S., Ren, B., and Iwase, S. (2017). LSD1/KDM1A Maintains Genome-wide Homeostasis of Transcriptional Enhancers. bioRxiv.

Alarcón, J.M., Malleret, G., Touzani, K., Vronskaya, S., Ishii, S., Kandel, E.R., and Barco, A. (2004). Chromatin Acetylation, Memory, and LTP Are Impaired in CBP+/− Mice: A Model for the Cognitive Deficit in Rubinstein-Taybi Syndrome and Its Amelioration. Neuron 42, 947–959.

Allen Institute for Brain Science (2015). Allen Developing Mouse Brain Atlas [Internet]. Available from http://developingmouse.brain-map.org.

Bateup, H.S., Johnson, C.A., Denefrio, C.L., Saulnier, J.L., Kornacker, K., and Sabatini, B.L. (2013). Excitatory/inhibitory synaptic imbalance leads to hippocampal hyperexcitability in mouse models of tuberous sclerosis. Neuron 78, 510–522.

Benevento, M., Iacono, G., Selten, M., Ba, W., Oudakker, A., Frega, M., Keller, J., Mancini, R., Lewerissa, E., Kleefstra, T., et al. (2016). Histone Methylation by the Kleefstra Syndrome Protein EHMT1 Mediates Homeostatic Synaptic Scaling. Neuron 91, 341–355.

Benito, E., and Barco, A. (2015). The neuronal activity-driven transcriptome. Mol Neurobiol 51, 1071–1088.

Bi, W., Ohyama, T., Nakamura, H., Yan, J., Visvanathan, J., Justice, M.J., and Lupski, J.R. (2005). Inactivation of Rai1 in mice recapitulates phenotypes observed in chromosome engineered mouse models for Smith-Magenis syndrome. Hum Mol Genet 14, 983–995.

Bi, W., Saifi, G.M., Shaw, C.J., Walz, K., Fonseca, P., Wilson, M., Potocki, L., and Lupski, J.R. (2004). Mutations of RAI1, a PHD-containing protein, in nondeletion patients with Smith-Magenis syndrome. Hum Genet 115, 515–524.

Bi, W., Yan, J., Shi, X., Yuva-Paylor, L.a., Antalffy, B.a., Goldman, A., Yoo, J.W., Noebels, J.L., Armstrong, D.L., Paylor, R., et al. (2007). Rai1 deficiency in mice causes learning impairment and motor dysfunction, whereas Rai1 heterozygous mice display minimal behavioral phenotypes. Hum Mol Genet 16, 1802–1813.

Boone, P.M., Reiter, R.J., Glaze, D.G., Tan, D.X., Lupski, J.R., and Potocki, L. (2011). Abnormal circadian rhythm of melatonin in Smith-Magenis syndrome patients with RAI1 point mutations. Am J Med Genet A 155A, 2024–2027.

Bourgeron, T. (2015). From the genetic architecture to synaptic plasticity in autism spectrum disorder. Nature Reviews Neuroscience 16, 551.

Bourtchouladze, R., Lidge, R., Catapano, R., Stanley, J., Gossweiler, S., Romashko, D., Scott, R., and Tully, T. (2003). A mouse model of Rubinstein-Taybi syndrome: Defective long-term memory is ameliorated by inhibitors of phosphodiesterase 4. Proceedings of the National Academy of Sciences 100, 10518.

Brakeman, P.R., Lanahan, A.A., O’Brien, R., Roche, K., Barnes, C.A., Huganir, R.L., and Worley, P.F. (1997). Homer: a protein that selectively binds metabotropic glutamate receptors. Nature 386, 284.

Bramham, C.R., Worley, P.F., Moore, M.J., and Guzowski, J.F. (2008). The Immediate Early Gene <em>Arc</em>/<em>Arg3.1</em>: Regulation, Mechanisms, and Function. The Journal of Neuroscience 28, 11760–11767.

Burns, B., Schmidt, K., Williams, S.R., Kim, S., Girirajan, S., and Elsea, S.H. (2010). Rai1 haploinsufficiency causes reduced Bdnf expression resulting in hyperphagia, obesity and altered fat distribution in mice and humans with no evidence of metabolic syndrome. Hum Mol Genet 19, 4026–4042.

Carmona-Mora, P., Canales, C.P., Cao, L., Perez, I.C., Srivastava, A.K., Young, J.I., and Walz, K. (2012). RAI1 Transcription Factor Activity Is Impaired in Mutants Associated with Smith-Magenis Syndrome. PLoS One 7.

Carmona-Mora, P., Encina, C.a., Canales, C.P., Cao, L., Molina, J., Kairath, P., Young, J.I., and Walz, K. (2010). Functional and cellular characterization of human Retinoic Acid Induced 1 (RAI1) mutations associated with Smith-Magenis Syndrome. BMC Mol Biol 11, 63–63.

Cleary, M.D., Meiering, C.D., Jan, E., Guymon, R., and Boothroyd, J.C. (2005). Biosynthetic labeling of RNA with uracil phosphoribosyltransferase allows cell-specific microarray analysis of mRNA synthesis and decay. Nat Biotechnol 23, 232.

Core, L.J., Waterfall, J.J., and Lis, J.T. (2008). Nascent RNA Sequencing Reveals Widespread Pausing and Divergent Initiation at Human Promoters. Science 322, 1845.

Darvekar, S., Johnsen, Sylvia S., Eriksen, Agnete B., Johansen, T., and Sjøttem, E. (2012). Identification of two independent nucleosome-binding domains in the transcriptional co-activator SPBP. Biochem J 442, 65–75.

Darvekar, S., Rekdal, C., Johansen, T., and Sjøttem, E. (2013). A Phylogenetic Study of SPBP and RAI1: Evolutionary Conservation of Chromatin Binding Modules. PLoS One 8, 1–11.

de Vivo, L., Bellesi, M., Marshall, W., Bushong, E.A., Ellisman, M.H., Tononi, G., and Cirelli, C. (2017). Ultrastructural evidence for synaptic scaling across the wake/sleep cycle. Science 355, 507.

Diering, G.H., Nirujogi, R.S., Roth, R.H., Worley, P.F., Pandey, A., and Huganir, R.L. (2017). Homer1a drives homeostatic scaling-down of excitatory synapses during sleep. Science 355, 511.

Ebert, D.H., and Greenberg, M.E. (2013). Activity-dependent neuronal signalling and autism spectrum disorder. Nature 493, 327–337.

Elsea, S.H., and Williams, S.R. (2011). Smith-Magenis syndrome: haploinsufficiency of RAI1 results in altered gene regulation in neurological and metabolic pathways. Expert Rev Mol Med 13, e14–e14.

Fernandes, D., and Carvalho, A.L. (2016). Mechanisms of homeostatic plasticity in the excitatory synapse. J Neurochem.

Fragoso, Y.D., Stoney, P.N., Shearer, K.D., Sementilli, A., Nanescu, S.E., Sementilli, P., and McCaffery, P. (2014). Expression in the human brain of retinoic acid induced 1, a protein associated with neurobehavioural disorders. Brain structure & function.

Girirajan, S., Elsas, L.J., Devriendt, K., and Elsea, S.H. (2005). RAI1 variations in Smith-Magenis syndrome patients without 17p11.2 deletions. J Med Genet 42, 820–828.

Girirajan, S., Truong, H.T., Blanchard, C.L., and Elsea, S.H. (2009). A functional network module for Smith-Magenis syndrome. Clin Genet 75, 364–374.

Glock, C., Heumüller, M., and Schuman, E.M. (2017). mRNA transport & local translation in neurons. Curr Opin Neurobiol 45, 169–177.

Gropman, A.L., Duncan, W.C., and Smith, A.C. (2006). Neurologic and developmental features of the Smith-Magenis syndrome (del 17p11.2). Pediatr Neurol 34, 337–350.

Guan, J.S., Haggarty, S.J., Giacometti, E., Dannenberg, J.H., Joseph, N., Gao, J., Nieland, T.J., Zhou, Y., Wang, X., Mazitschek, R., et al. (2009). HDAC2 negatively regulates memory formation and synaptic plasticity. Nature 459, 55–60.

Gupta-Agarwal, S., Franklin, A.V., Deramus, T., Wheelock, M., Davis, R.L., McMahon, L.L., and Lubin, F.D. (2012). G9a/GLP histone lysine dimethyltransferase complex activity in the hippocampus and the entorhinal cortex is required for gene activation and silencing during memory consolidation. J Neurosci 32, 5440–5453.

Gupta-Agarwal, S., Jarome, T.J., Fernandez, J., and Lubin, F.D. (2014). NMDA receptor- and ERK-dependent histone methylation changes in the lateral amygdala bidirectionally regulate fear memory formation. Learn Mem 21, 351–362.

Guzman-Karlsson, M.C., Meadows, J.P., Gavin, C.F., Hablitz, J.J., and Sweatt, J.D. (2014). Transcriptional and epigenetic regulation of Hebbian and non-Hebbian plasticity. Neuropharmacology 80, 3–17.

Hasel, P., Dando, O., Jiwaji, Z., Baxter, P., Todd, A.C., Heron, S., Markus, N.M., McQueen, J., Hampton, D.W., Torvell, M., et al. (2017). Neurons and neuronal activity control gene expression in astrocytes to regulate their development and metabolism. Nat Commun 8, 15132.

Hermey, G., Blüthgen, N., and Kuhl, D. (2017). Neuronal activity-regulated alternative mRNA splicing. The International Journal of Biochemistry & Cell Biology 91, 184–193.

Hrvatin, S., Hochbaum, D.R., Nagy, M.A., Cicconet, M., Robertson, K., Cheadle, L., Zilionis, R., Ratner, A., Borges-Monroy, R., Klein, A.M., et al. (2018). Single-cell analysis of experience-dependent transcriptomic states in the mouse visual cortex. Nat Neurosci 21, 120–129.

Huang, W.-H., Wang, D.C., Allen, W.E., Klope, M., Hu, H., Shamloo, M., and Luo, L. (2018). Early adolescent Rai1 reactivation reverses transcriptional and social interaction deficits in a mouse model of Smith–Magenis syndrome. Proceedings of the National Academy of Sciences 115, 10744.

Huang, W.H., Guenthner, C.J., Xu, J., Nguyen, T., Schwarz, L.A., Wilkinson, A.W., Gozani, O., Chang, H.Y., Shamloo, M., and Luo, L. (2016). Molecular and Neural Functions of Rai1, the Causal Gene for Smith-Magenis Syndrome. Neuron 92, 392–406.

Ibata, K., Sun, Q., and Turrigiano, G.G. (2008). Rapid synaptic scaling induced by changes in postsynaptic firing. Neuron 57, 819–826.

Igaz, L.M., Vianna, M.R.M., Medina, J.H., and Izquierdo, I. (2002). Two Time Periods of Hippocampal mRNA Synthesis Are Required for Memory Consolidation of Fear-Motivated Learning. The Journal of Neuroscience 22, 6781.

Ito, S., D’Alessio, A.C., Taranova, O.V., Hong, K., Sowers, L.C., and Zhang, Y. (2010). Role of Tet proteins in 5mC to 5hmC conversion, ES-cell self-renewal and inner cell mass specification. Nature 466, 1129.

Iwase, S., Brookes, E., Agarwal, S., Badeaux, A.I., Ito, H., Vallianatos, C.N., Tomassy, G.S., Kasza, T., Lin, G., Thompson, A., et al. (2016). A Mouse Model of X-linked Intellectual Disability Associated with Impaired Removal of Histone Methylation. Cell Rep 14, 1000–1009.

Kerimoglu, C., Agis-Balboa, R.C., Kranz, A., Stilling, R., Bahari-Javan, S., Benito-Garagorri, E., Halder, R., Burkhardt, S., Stewart, A.F., and Fischer, A. (2013). HistoneMethyltransferase MLL2 (KMT2B) Is Required for Memory Formation in Mice. The Journal of Neuroscience 33, 3452.

Kim, D., Pertea, G., Trapnell, C., Pimentel, H., Kelley, R., and Salzberg, S.L. (2013). TopHat2: accurate alignment of transcriptomes in the presence of insertions, deletions and gene fusions. Genome Biol 14, R36.

Kim, J.H., Karnovsky, A., Mahavisno, V., Weymouth, T., Pande, M., Dolinoy, D.C., Rozek, L.S., and Sartor, M.A. (2012). LRpath analysis reveals common pathways dysregulated via DNA methylation across cancer types. BMC Genomics 13, 526.

Kuhn, M., Wolf, E., Maier, J.G., Mainberger, F., Feige, B., Schmid, H., Bürklin, J., Maywald, S., Mall, V., Jung, N.H., et al. (2016). Sleep recalibrates homeostatic and associative synaptic plasticity in the human cortex. Nature Communications 7, 12455.

Lacaria, M., Gu, W., and Lupski, J.R. (2013). Circadian abnormalities in mouse models of smith-magenis syndrome: Evidence for involvement of RAI1. American Journal of Medical Genetics, Part A 161, 1561–1568.

Langmead, B., and Salzberg, S.L. (2012). Fast gapped-read alignment with Bowtie 2. Nat Methods 9, 357.

Lee, C., Patil, S., and Sartor, M.A. (2016). RNA-Enrich: a cut-off free functional enrichment testing method for RNA-seq with improved detection power. Bioinformatics 32, 1100–1102.

Liao, Y., Smyth, G.K., and Shi, W. (2014). featureCounts: an efficient general purpose program for assigning sequence reads to genomic features. Bioinformatics 30, 923–930.

Lim, C.-S., Nam, H.J., Lee, J., Kim, D., Choi, J.E., Kang, S.J., Kim, S., Kim, H., Kwak, C., Shim, K.-W., et al. (2017). PKCα-mediated phosphorylation of LSD1 is required for presynaptic plasticity and hippocampal learning and memory. Sci Rep 7, 4912.

Love, M.I., Huber, W., and Anders, S. (2014). Moderated estimation of fold change and dispersion for RNA-seq data with DESeq2. Genome Biol 15, 550.

Mao, W., Salzberg, A.C., Uchigashima, M., Hasegawa, Y., Hock, H., Watanabe, M., Akbarian, S., Kawasawa, Y.I., and Futai, K. (2018). Activity-Induced Regulation of Synaptic Strength through the Chromatin Reader L3mbtl1. Cell Reports 23, 3209–3222.

Miller, K.D., and MacKay, D.J.C. (1994). The Role of Constraints in Hebbian Learning. Neural Comput 6, 100–126.

Molina, J., Carmona-Mora, P., Chrast, J., Krall, P.M., Canales, C.P., Lupski, J.R., Reymond, A., and Walz, K. (2008). Abnormal social behaviors and altered gene expression rates in a mouse model for Potocki-Lupski syndrome. Hum Mol Genet 17, 2486–2495.

Mullins, C., Fishell, G., and Tsien, R.W. (2016). Unifying Views of Autism Spectrum Disorders: A Consideration of Autoregulatory Feedback Loops. Neuron 89, 1131–1156.

Neelamegam, R., Ricq, E.L., Malvaez, M., Patnaik, D., Norton, S., Carlin, S.M., Hill, I.T., Wood, M.A., Haggarty, S.J., and Hooker, J.M. (2012). Brain-penetrant LSD1 inhibitors can block memory consolidation. ACS Chem Neurosci 3, 120–128.

Oike, Y., Hata, A., Mamiya, T., Kaname, T., Noda, Y., Suzuki, M., Yasue, H., Nabeshima, T., Araki, K., and Yamamura, K.-i. (1999). Truncated CBP Protein Leads to Classical Rubinstein—Taybi Syndrome Phenotypes in Mice: Implications for a Dominant-Negative Mechanism. Hum Mol Genet 8, 387–396.

Okuno, H. (2011). Regulation and function of immediate-early genes in the brain: Beyond neuronal activity markers. Neurosci Res 69, 175–186.

Oliveira, A.M.M. (2016). DNA methylation: a permissive mark in memory formation and maintenance. Learn Memory 23, 587–593.

Paulsen, M.T., Veloso, A., Prasad, J., Bedi, K., Ljungman, E.A., Magnuson, B., Wilson, T.E., and Ljungman, M. (2014). Use of Bru-Seq and BruChase-Seq for genome-wide assessment of the synthesis and stability of RNA. Methods (San Diego, Calif) 67, 45–54.

Paulsen, M.T., Veloso, A., Prasad, J., Bedi, K., Ljungman, E.A., Tsan, Y.-C., Chang, C.-W., Tarrier, B., Washburn, J.G., Lyons, R., et al. (2013). Coordinated regulation of synthesis and stability of RNA during the acute TNF-induced proinflammatory response. Proceedings of the National Academy of Sciences 110, 2240–2245.

Potocki, L., Bi, W., Treadwell-Deering, D., Carvalho, C.M., Eifert, A., Friedman, E.M., Glaze, D., Krull, K., Lee, J.A., Lewis, R.A., et al. (2007). Characterization of PotockiLupski syndrome (dup(17)(p11.2p11.2)) and delineation of a dosage-sensitive critical interval that can convey an autism phenotype. Am J Hum Genet 80, 633–649.

Potocki, L., Glaze, D., Tan, D., Park, S., Kashork, C., Shaffer, L., Reiter, R., and Lupski, J. (2000). Circadian rhythm abnormalities of melatonin in Smith-Magenis syndrome. J Med Genet 37, 428–433.

Risher, W.C., and Eroglu, C. (2012). Thrombospondins as key regulators of synaptogenesis in the central nervous system. Matrix Biol 31, 170–177.

Rudenko, A., Dawlaty, M.M., Seo, J., Cheng, A.W., Meng, J., Le, T., Faull, K.F., Jaenisch, R., and Tsai, L.H. (2013). Tet1 is critical for neuronal activity-regulated gene expression and memory extinction. Neuron 79, 1109–1122.

Schanzenbächer, Christoph T., Sambandan, S., Langer, Julian D., and Schuman, Erin M. (2016). Nascent Proteome Remodeling following Homeostatic Scaling at Hippocampal Synapses. Neuron 92, 358–371.

Schaukowitch, K., Reese, A.L., Kim, S.K., Kilaru, G., Joo, J.Y., Kavalali, E.T., and Kim, T.K. (2017). An Intrinsic Transcriptional Program Underlying Synaptic Scaling during Activity Suppression. Cell Rep 18, 1512–1526.

Slager, R.E., Newton, T.L., Vlangos, C.N., Finucane, B., and Elsea, S.H. (2003). Mutations in RAI1 associated with Smith–Magenis syndrome. Nat Genet 33, 466.

Soden, M.E., and Chen, L. (2010). Fragile X protein FMRP is required for homeostatic plasticity and regulation of synaptic strength by retinoic acid. J Neurosci 30, 16910–16921.

Supek, F., Bošnjak, M., Škunca, N., and Šmuc, T. (2011). REVIGO Summarizes and Visualizes Long Lists of Gene Ontology Terms. PLoS One 6, e21800.

Tachibana, M., Ueda, J., Fukuda, M., Takeda, N., Ohta, T., Iwanari, H., Sakihama, T., Kodama, T., Hamakubo, T., and Shinkai, Y. (2005). Histone methyltransferases G9a and GLP form heteromeric complexes and are both crucial for methylation of euchromatin at H3-K9. Genes Dev 19, 815–826.

Tasic, B., Menon, V., Nguyen, T.N., Kim, T.K., Jarsky, T., Yao, Z., Levi, B., Gray, L.T., Sorensen, S.A., Dolbeare, T., et al. (2016). Adult mouse cortical cell taxonomy revealed by single cell transcriptomics. Nat Neurosci 19, 335.

Turrigiano, G.G. (2008). The self-tuning neuron: synaptic scaling of excitatory synapses. Cell 135, 422–435.

Turrigiano, G.G. (2017). The dialectic of Hebb and homeostasis. Philosophical Transactions of the Royal Society B: Biological Sciences 372.

Vallianatos, C.N., Farrehi, C., Friez, M.J., Burmeister, M., Keegan, C.E., and Iwase, S. (2018). Altered Gene-Regulatory Function of KDM5C by a Novel Mutation Associated With Autism and Intellectual Disability. Front Mol Neurosci 11.

Vogel-Ciernia, A., Matheos, D.P., Barrett, R.M., Kramar, E.A., Azzawi, S., Chen, Y., Magnan, C.N., Zeller, M., Sylvain, A., Haettig, J., et al. (2013). The neuron-specific chromatin regulatory subunit BAF53b is necessary for synaptic plasticity and memory. Nat Neurosci 16, 552–561.

Wang, J., Telese, F., Tan, Y., Li, W., Jin, C., He, X., Basnet, H., Ma, Q., Merkurjev, D., Zhu, X., et al. (2015). LSD1n is an H4K20 demethylase regulating memory formation via transcriptional elongation control. Nat Neurosci 18, 1256–1264.

West, A.E., Griffith, E.C., and Greenberg, M.E. (2002). Regulation of transcription factors by neuronal activity. Nat Rev Neurosci 3, 921–931.

Widagdo, J., and Anggono, V. (2018). The m6A-epitranscriptomic signature in neurobiology: from neurodevelopment to brain plasticity. J Neurochem 0.

Williams, S.R., Zies, D., Mullegama, S.V., Grotewiel, M.S., and Elsea, S.H. (2012). Smith-magenis syndrome results in disruption of CLOCK gene transcription and reveals an integral role for RAI1 in the maintenance of circadian rhythmicity. Am J Hum Genet 90, 941–949.

Yee, A.X., Hsu, Y.-T., and Chen, L. (2017). A metaplasticity view of the interaction between homeostatic and Hebbian plasticity. Philosophical Transactions of the Royal Society B: Biological Sciences 372.

Yu, H., Su, Y., Shin, J., Zhong, C., Guo, J.U., Weng, Y.-l., Gao, F., Geschwind, D.H., Coppola, G., Ming, G.-l., et al. (2015). Tet3 regulates synaptic transmission and homeostatic plasticity via DNA oxidation and repair. 18, 7–9.

Zajaczkowski, E.L., Zhao, Q.-Y., Zhang, Z.H., Li, X., Wei, W., Marshall, P.R., Leighton, L.J., Nainar, S., Feng, C., Spitale, R.C., et al. (2018). Bioorthogonal Metabolic Labeling of Nascent RNA in Neurons Improves the Sensitivity of Transcriptome-Wide Profiling. ACS Chem Neurosci 9, 1858–1865.

Zhang, Y., Chen, K., Sloan, S.A., Bennett, M.L., Scholze, A.R., O’Keeffe, S., Phatnani, H.P., Guarnieri, P., Caneda, C., Ruderisch, N., et al. (2014). An RNA-sequencing transcriptome and splicing database of glia, neurons, and vascular cells of the cerebral cortex. J Neurosci 34, 11929–11947.

Zhang, Y., Liu, T., Meyer, C.A., Eeckhoute, J., Johnson, D.S., Bernstein, B.E., Nusbaum, C., Myers, R.M., Brown, M., Li, W., et al. (2008). Model-based analysis of ChIP-Seq (MACS). Genome Biol 9, R137.

Zhong, X., Li, H., and Chang, Q. (2012). MeCP2 phosphorylation is required for modulating synaptic scaling through mGluR5. J Neurosci 32, 12841–12847.

